# IRF7 controls spontaneous autoimmune germinal center and plasma cell checkpoints

**DOI:** 10.1101/2025.02.04.636277

**Authors:** Adam J. Fike, Kristen N. Bricker, Michael V. Gonzalez, Anju Maharjan, Tien Bui, Keomonyroth Nuon, Scott M. Emrich, Julia L. Weber, Sara A. Luckenbill, Nicholas M. Choi, Renan Sauteraud, Dajiang J. Liu, Nancy J. Olsen, Roberto Caricchio, Mohamed Trebak, Sathi Babu Chodisetti, Ziaur S.M. Rahman

## Abstract

How IRF7 promotes autoimmune B cell responses and systemic autoimmunity is unclear. Analysis of spontaneous SLE-prone mice deficient in IRF7 uncovered the IRF7 role in regulating autoimmune germinal center (GC), plasma cell (PC) and autoantibody responses and disease. IRF7, however, was dispensable for foreign antigen driven GC, PC and antibody responses. Competitive bone marrow (BM) chimeras highlighted the importance of IRF7 in hematopoietic cells in spontaneous GC and PC differentiation. Single-cell-RNAseq of SLE-prone B cells indicated IRF7 mediated B cell differentiation through GC and PC fates. Mechanistic studies revealed that IRF7 promoted B cell differentiation through GC and PC fates by regulating the transcriptome, translation, and metabolism of SLE-prone B cells. Mixed BM chimeras demonstrated a requirement for B cell-intrinsic IRF7 in IgG autoantibody production but not sufficient for promoting spontaneous GC and PC responses. Altogether, we delineate previously unknown B cell-intrinsic and -extrinsic mechanisms of IRF7-promoted spontaneous GC and PC responses, loss of tolerance, autoantibody production and SLE development.

**Summary:** Fike et al. describe previously unknown mechanisms by which IRF7 controls autoimmune B cell and autoantibody responses. Mechanistic studies guided by single-cell-RNAseq and ChIPseq reveal that IRF7 promotes B cell differentiation through germinal center and plasma cell fates by regulating the transcriptome, translation, and metabolism of lupus-prone B cells.

## INTRODUCTION

Systemic lupus erythematosus (SLE) is a debilitating autoimmune disease initiated by dysregulated B cell responses and the emergence of autoreactive B cells (Karrar and Cunninghame Graham, 2018; Nashi et al., 2010; Rawlings et al., 2017). Autoreactive B cells in SLE produce antinuclear antibodies (ANA) that deposit as immune complexes in various tissues including kidneys and vasculature, causing inflammation and end organ disease (Bagavant and Fu, 2009). Genetic predisposition in concert with toll-like receptor (TLR) activation by self-nucleic acid permits B cell escape of tolerance mechanisms, leading to the emergence of autoreactive B cells (Celhar and Fairhurst, 2014; Christensen and Shlomchik, 2007; Green et al., 2009). TLR signaling promotes aberrant differentiation of autoreactive B cells into plasma (PC) and geminal center (GC) cells to generate autoantibody producing B cells (Brown et al., 2022a; Christensen et al., 2006; Deane et al., 2007; Jackson et al., 2014; Soni et al., 2014); however, the signals downstream TLR activation and mechanisms that regulate autoreactive B cells in SLE remain unclear.

We and others previously demonstrated a B cell intrinsic requirement for TLR7 in promoting autoimmune GC and PC responses and systemic autoimmunity (Brown et al., 2022a; Christensen et al., 2006; Cosgrove et al., 2023; Deane et al., 2007; Hwang et al., 2012; Jackson et al., 2014; Soni et al., 2014). Stimulation of endosomal TLRs by nucleic acids including TLR7 activates NFκB family members, and interferon regulatory factors 5 (IRF5) and 7 (IRF7). To date, numerous studies have characterized the involvement of IRF5 and IRF5 risk variants in SLE in mice and humans (Ban et al., 2021; Banga et al., 2020; Cham et al., 2012; Pellerin et al., 2021; Richez et al., 2010). IRF7 was previously shown to promote autoantibody responses but not lupus nephritis in a chemically (pristane)-induced B6 model of SLE (Miyagawa et al., 2016). IRF7 risk alleles are also implicated in the development of human SLE (Salloum et al., 2010; Yasuda et al., 2007). However, IRF7-expressing cell type(s) and mechanisms by which IRF7 drives autoreactive B cell differentiation through PC and GC checkpoints, leading to autoantibody-secreting PCs and autoantibody production in SLE remain unknown. IRF7 has long been thought to contribute to SLE by regulating type 1 interferon (T1-IFN) production by plasmacytoid dendritic cells (pDCs) (Honda et al., 2005; Yasuda et al., 2007), although IRF7 is also expressed in B cells and monocytes (Ning et al., 2011). Whether and how IRF7 may promote autoimmune GC and PC responses, autoantibody production and disease in spontaneous SLE models via functioning in cell types other than pDCs is unknown. It is also unclear whether IRF7 transcriptionally controls SLE-promoting gene programs in B cells other than T1-IFN genes.

Here, using FcγRIIB^−/−^ and FcγRIIB^−/−^Yaa spontaneous SLE mouse models, we identified a major role for IRF7 in promoting spontaneous autoimmune PC, GC and autoantibody responses and SLE-like disease but not in foreign-antigen-driven PC, GC, and antibody responses. Single-cell-RNAseq (scRNAseq) of B cells from FcγRIIB^−/−^ IRF7^−/−^ and FcγRIIB^−/−^ control mice indicated a role for IRF7-driven genetic programs in regulating B cell differentiation through pre-GC to GC and PC populations. Mixed bone marrow competition chimeras demonstrated a role for IRF7 expression in hematopoietic compartments in driving GC and PC but not myeloid and T cell responses in FcγRIIB^−/−^ mice. Using B cell-specific mixed bone marrow chimeras we further demonstrated a B cell-intrinsic requirement of IRF7 in the production of class-switched IgG autoantibodies but not for promoting overall spontaneous autoimmune PC and GC responses. Together, we delineate B cell-intrinsic and -extrinsic functions for IRF7 in regulating B cell transcription, translation, and metabolism, and in breaching tolerance and promoting autoimmune GC and PC responses and systemic autoimmunity development.

## RESULTS

### IRF7 promotes autoimmune GC and PC responses in spontaneous SLE models

To identify the mechanisms by which IRF7 regulates autoimmune GC and PC responses and systemic autoimmunity, we first crossed IRF7-deficient (IRF7^−/−^) mice to well-characterized SLE-prone FcγRIIB^−/−^ mice (Bolland et al., 2002; Richez et al., 2010; Shinde et al., 2018). FcγRIIB^−/−^ mice, originally generated in 129 background and subsequently backcrossed to B6 mice, are deficient in FcγRIIB and carry 129-derived signaling lymphocyte activation molecule (SLAM) family genes that are tightly linked to the *Fcgr2b* gene (Soni et al., 2015). FcγRIIB^−/−^ mice promote many of the manifestations of human SLE including dysregulated GC and PC responses, antinuclear antibodies (ANAs), and renal disease (Bolland et al., 2002; Shinde et al., 2018; Soni et al., 2015). FcγRIIB^−/−^ mice deficient in IRF7 (FcγRIIB^−/−^IRF7^−/−^) had reduced GC, effector T cell (Teff) and T follicular helper (Tfh) cell, and PC (Figure 1A-H) responses at 2, 4 and 6 mo ages. FcγRIIB^−/−^IRF7^−/−^ mice had reduced numbers of autoantibody-producing AFCs in the spleen and bone marrow (Figure 1I, J) and dampened serum ANA reactivity (Figure 1K).

**Figure 1:**
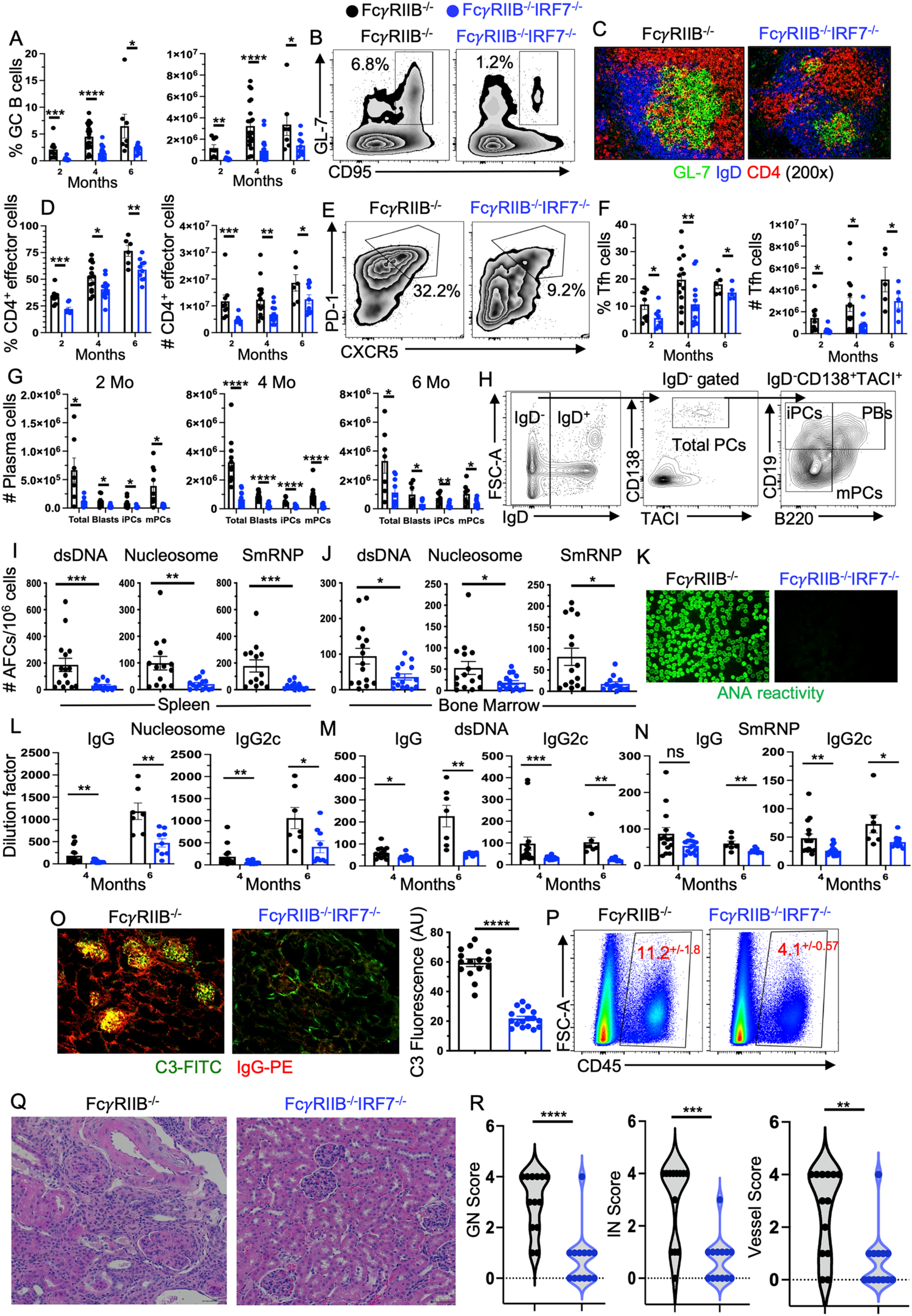
IRF7 promotes autoimmunity in the FcγRIIB^−/−^ mouse model of SLE. (A-H) B and T cell responses were evaluated in the spleens of FcγRIIB^−/−^ and FcγRIIB^−/−^IRF7^−/−^ mice at 2, 4 and 6 mo time points. (A) Frequency and number of B220^+^GL-7^+^CD95^+^ GC B cells of total B220^+^ B cells. (B) Representative flow plots showing the gating strategy of GC B cells. (C) Representative images showing IgD^−^GL-7^+^ GC B cells and CD4^+^ Tfh cells within the GC by staining splenic sections with anti-IgD (blue), GL-7 (green) and anti-CD4 (red). (D) Frequency and number of CD4^+^CD44^+^CD62L^−^ effector T cells of total CD4^+^ cells. (E) Flow cytometry gating strategy, and (F) frequency and number of CD4^+^CD44^+^CD62L^−^CXCR5^+^PD-1^+^ Tfh of total CD4^+^ cells. (G) Number and (H) flow cytometry gating strategy of CD138^+^TACI^+^ total plasma cells gated on IgD^−^ cells followed by gating on CD138^+^TACI^+^ total plasma cells for B220 and CD19 expression to identify B220^+^CD19^+^ plasmablast (PB), B220^−^CD19^+^ immature (iPC) and B220^−^ CD19^−^ mature (mPC) plasma cells based on B220 and CD19 expression. ELISpot assay quantifying the number of dsDNA-, nucleosome- and SmRNP-specific (I) splenic and (J) bone marrow AFCs. (K) Hep-2 immunofluorescent (IF) images showing ANA seropositivity. (L) Nucleosome-, (M) dsDNA- and (N) SmRNP-specific serum IgG and IgG2c titers. (O) IF images of kidney sections showing C3^+^ (green) and IgG^+^ (red) immune complex deposition. (P) Flow cytometry plots showing the gating strategy and frequency of CD45^+^ immune cells in the kidney. (Q) Representative images of kidney sections stained with periodic acid–Schiff (PAS) and (R) the pathology score for glomerulonephritis (GN), interstitial nephritis (IN) and vessels. Each symbol (panels A to P and R) represents an individual mouse (n = 7-22 mice per group) and data are presented as means ± SEM. Data in panels A to R represent two-four experiments per time point. P values were calculated via two-way Anova with Sidak multiple comparisons test (A-G) or an unpaired Student’s t-test (I-R) (*, p <0.05, **, p <0.01, ***, p <0.001, ****, p <0.0001).

In FcγRIIB^−/−^ mice, autoantibodies started accumulating at 2 months of age when only nucleosome-specific serum IgM, IgG and IgG2c titers were reduced in IRF7-deficient mice (Figure S1). Nucleosome-, dsDNA- and SmRNP-specific IgG and IgG2c titers were significantly reduced in FcγRIIB^−/−^IRF7^−/−^ mice at 4 and 6 months of age when substantial amounts of autoantibodies accumulated in FcγRIIB^−/−^ mice (Figure 1L-N). We also found reduced numbers of IgG1- and IgG2c-producing but not total IgG- and IgG3-producing splenic and bone marrow AFCs in FcγRIIB^−/−^IRF7^−/−^ mice compared to FcγRIIB^−/−^ control mice (Figure S2A, B). In addition, total serum IgG, IgG1, IgG2c and IgG3 titers were lower in FcγRIIB^−/−^IRF7^−/−^ mice than FcγRIIB^−/−^ control mice (Figure S2C). Immune complex (IC) deposition and immune cell infiltration in the kidney (Figure 1O, P) and kidney pathology (Figure 1Q) and scoring for glomerulonephritis (GN), interstitial nephritis (IN) and vessels (Figure 1R) were significantly reduced in FcγRIIB^−/−^IRF7^−/−^ mice. FcγRIIB^−/−^IRF7^−/−^ mice also had reduced myeloid cell populations including conventional (cDCs) and monocyte derived DCs (moDCs), although frequency and number of plasmacytoid DCs (pDCs) were increased (Figure S3).

Splenomegaly (Figure 2A), GC, and PC responses (Figure 2B-E), and age associated B cell (ABC) responses as well as Teff and Tfh cell responses (Figure 2G, H) were also reduced in IRF7-deficient FcγRIIB^−/−^yaa male mice (Figure 2F) that express two copies of TLR7 and present with robust and rapid systemic autoimmune responses and disease (Bolland et al., 2002; Richez et al., 2010). Additionally, FcγRIIB^−/−^yaaIRF7^−/−^ mice showed reduced numbers of autoantibody-producing splenic and bone marrow AFCs (Figure S4A, B) and reduced serum titers of IgG and IgG2c autoantibodies to RNA-associated antigen, SmRNP (Figure 2I). However, no significant changes were observed in myeloid cell responses except for a consistent increase in the number of pDCs in the absence of IRF7 (Figure S4C, D). High-dimensional flow cytometry identified 20 immune cell populations (Table 1) within the kidneys of these mice (Table 1, Figure 2J, K). IRF7 deficiency dampened immune cell infiltration into the kidney (Figure 2J, K) with significant effects on CD4^+^ T cells (population 1), DCs (population 9), and macrophages (population 13). In addition, FcγRIIB^−/−^yaaIRF7^−/−^ mice had reduced serum blood urea nitrogen (BUN) and creatinine levels (Figure 2L, M), indicators of improved renal function. Finally, FcγRIIB^−/−^yaaIRF7^−/−^ mice had improved survival (Figure 2N).

**Figure 2:**
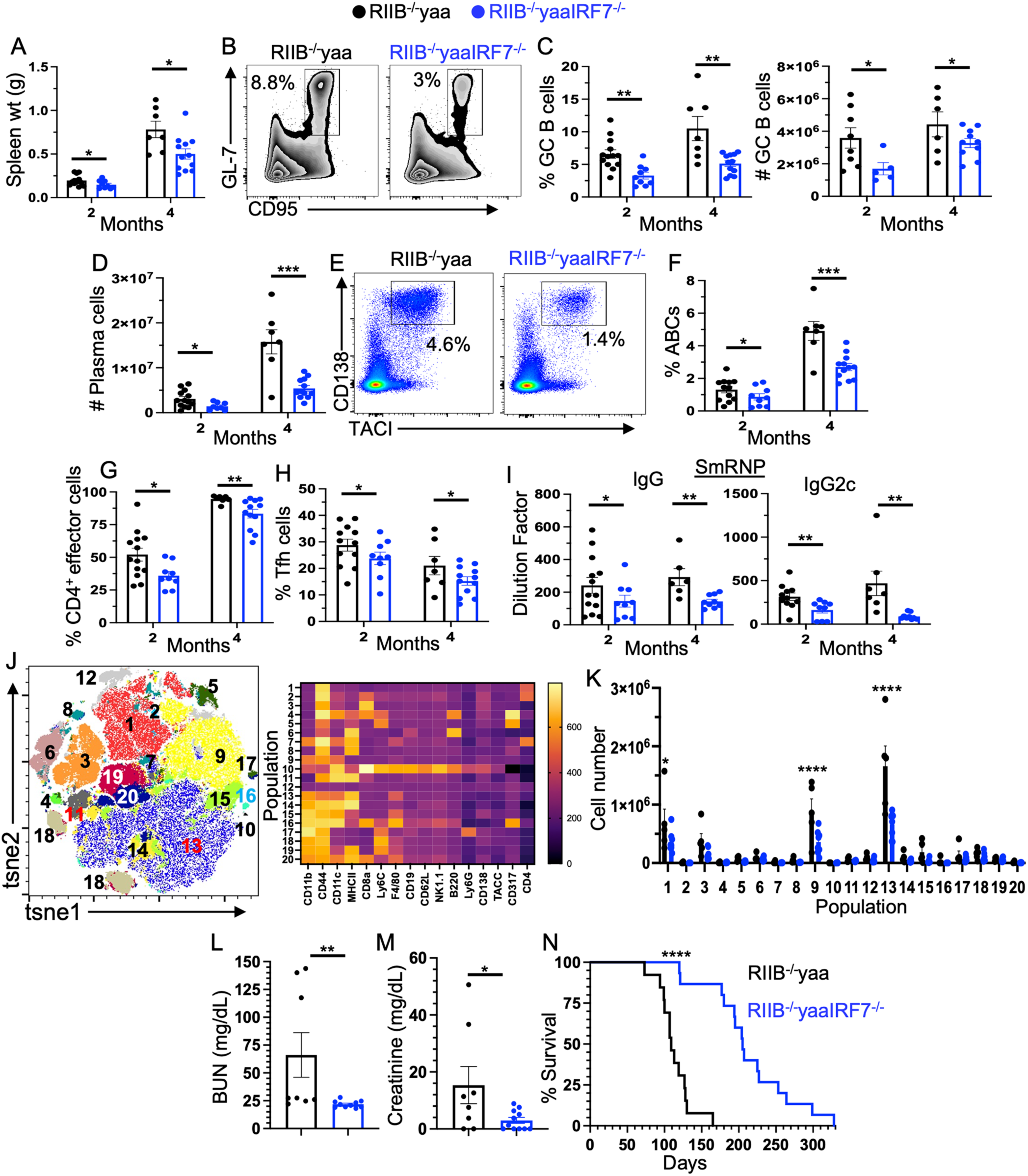
The major contribution of IRF7 to a TLR7-driven FcγRIIB^−/−^ model of SLE. (A-H) B and T cell responses were evaluated in the spleens of FcγRIIB^−/−^yaa (RIIB^−/−^yaa) and FcγRIIB^−/−^ yaaIRF7^−/−^ (RIIB^−/−^yaaIRF7^−/−^) mice at 2 and 4 mo age. (A) Spleen weight of RIIB^−/−^yaa and RIIB^−/−^ yaaIRF7^−/−^ mice. (B) Flow cytometry gating strategy, and (C) frequency and number of B220^+^GL-7^+^CD95^+^ GC B cells of total B220^+^ B cells. (D) Number and (E) flow cytometry gating strategy and frequency of CD138^+^TACI^+^ plasma cells in IgD^−^ B cells. (F) Frequency of IgD^−^ CD11b^+^CD11c^+^ age associated B cells (ABCs) of B220^+^CD19^+^ B cells in the spleen. Frequency of (G) CD4^+^CD44^+^CD62L^−^ effector T cells of total CD4^+^ cells and (H) CD4^+^CD44^+^CD62L^−^ CXCR5^+^PD-1^+^ Tfh of total CD4^+^ cells. (I) Serum IgG and IgG2c anti-SmRNP antibodies were measured by ELISA. (J, K) tsne plot generated from high dimensional flow cytometry analysis of kidney immune cell infiltrates identifying 20 populations. (L) BUN and (M) creatinine levels were measured in sera collected from 4 mo old mice. (N) The percentage of survival in RIIB^−/−^yaa and RIIB^−/−^yaaIRF7^−/−^ mice. Each symbol represents an individual mouse (n = 7-13 mice per group) and data are presented as means ± SEM. Data in each panel represent three experiments per time point. P values were calculated via two-way Anova with Dunn-Sidak correction (A-K), an unpaired Student’s t-test (L), Mann-Whitney test (M), or Log-rank (Mantel-Cox) test (N) (*, p <0.05, **, p <0.01, ***, p <0.001, ****, p <0.0001).

**Table 1:**
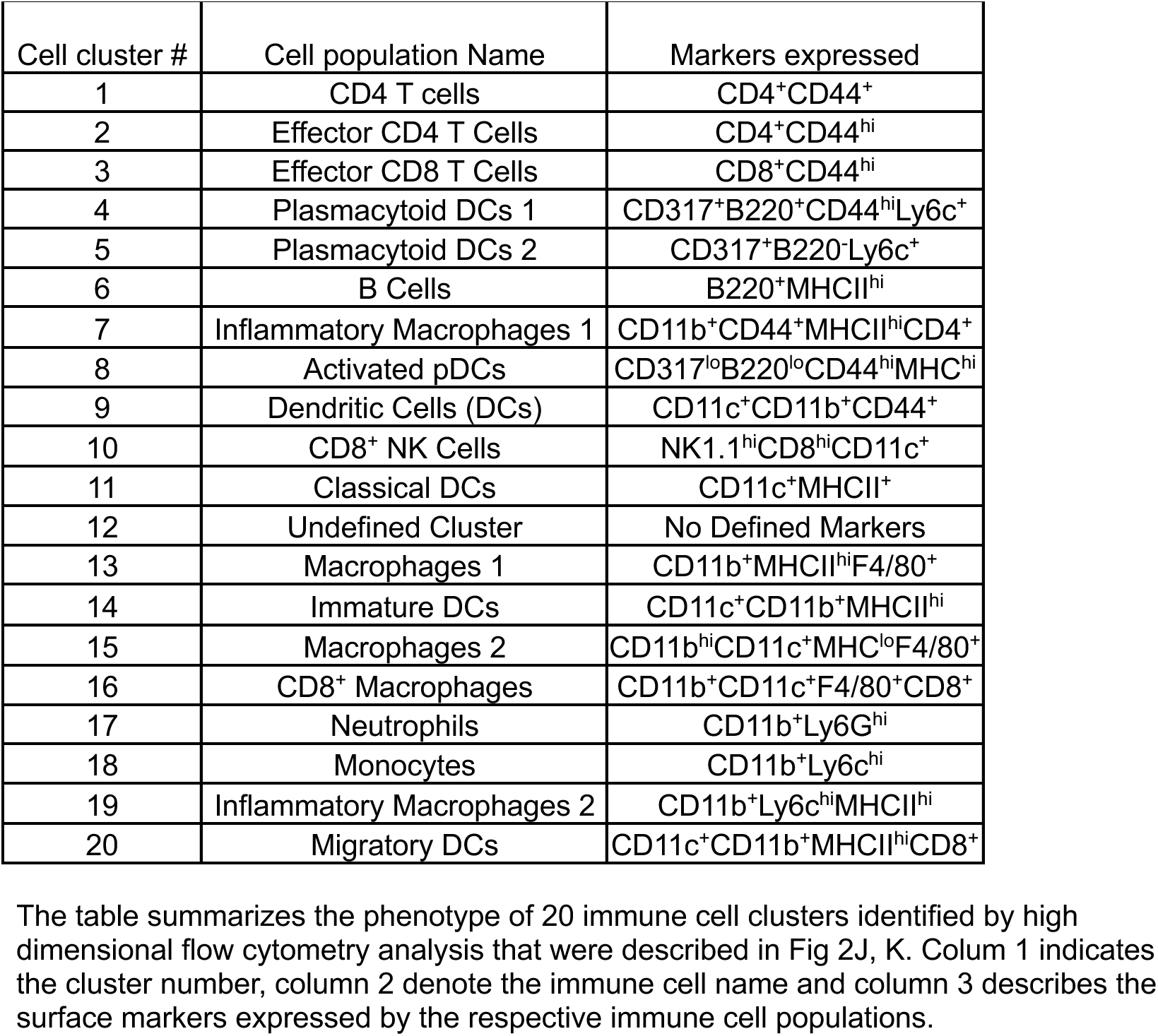
Phenotyping of kidney infiltrating immune cells by high-dimensional flow cytometric analysis.

Together, combined data from two SLE mouse models reinforce the role of IRF7 in controlling spontaneous autoimmune GC and PC responses, leading to autoantibody production and SLE-like disease.

### IRF7 is dispensable for foreign antigen driven GC and PC responses

We next determined the role of IRF7 in foreign antigen-driven GC, PC and antibody responses by immunizing B6 control and IRF7^−/−^ mice with NP-KLH in complete freund’s adjuvant (CFA). We observed no differences in the frequencies of total and NP-specific B cell, and GC B cell responses between the strains on 14d post-NP-KLH immunization (Figure 3A-D). Total and NP-specific CD138^+^TACI^+^ PC responses were similar between the strains except for an increased frequency of immature PCs in IRF7-deficient mice (Figure 3E-H). IRF7^−/−^ mice had reduced percentage of effector T cells, although no difference in Tfh frequency between the strains was observed (Figure 3I, J). We also observed no differences in high (NP_4_) and low (NP_28_) affinity NP-specific IgG and IgG1 Ab responses (Figure 3K, L).

**Figure 3.**
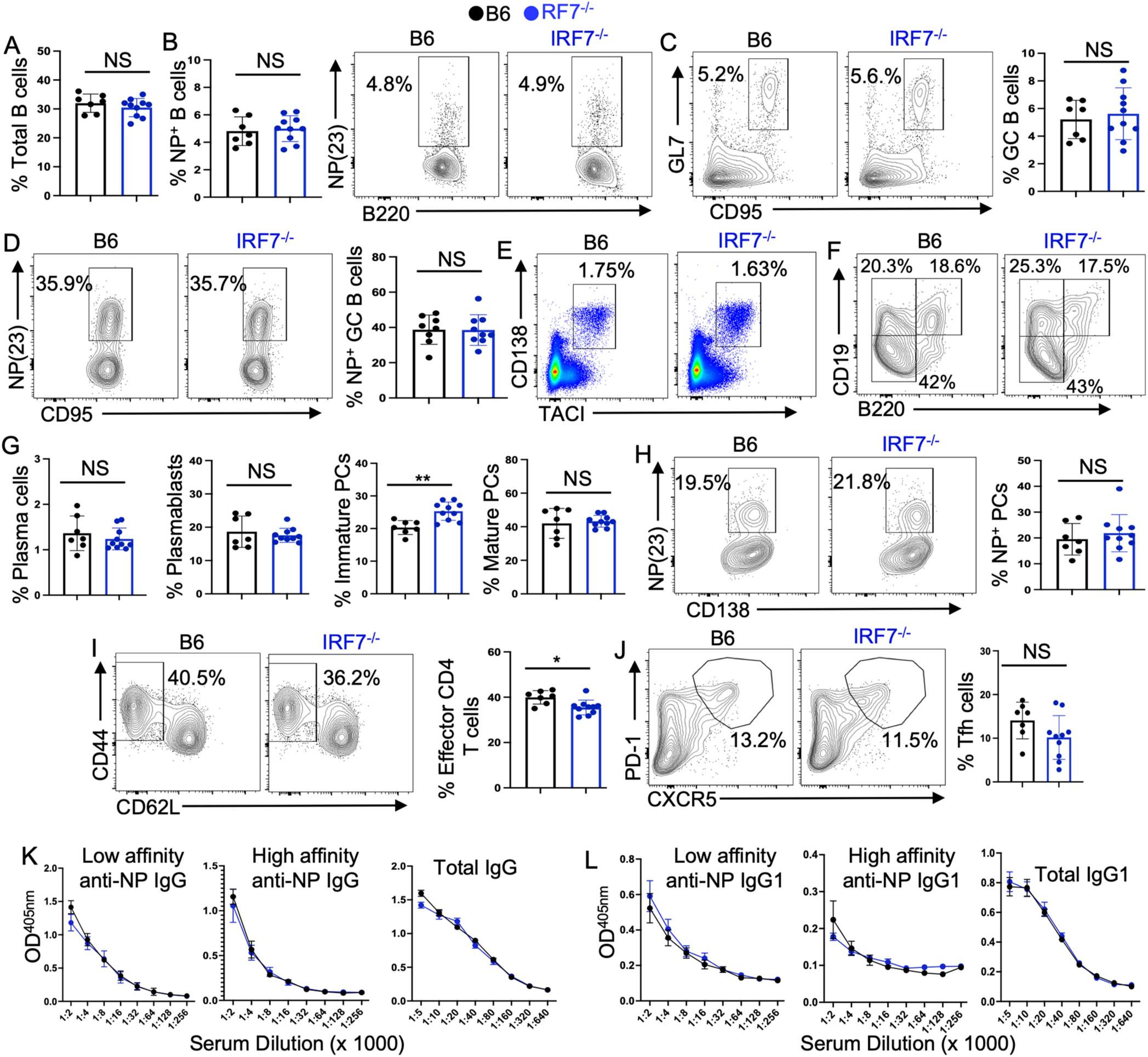
IRF7 signaling is not required for foreign antigen driven GC, PC and Ab responses. IRF7-deficient (IRF7^−/−^, blue circles) and C57BL/6 (B6, black circles) control mice were immunized with NP-KLH in CFA as described in Materials and Methods. The splenic GC, PC and antibody responses were analyzed on 14d post-immunization. Frequencies of (A) B220^+^ total and (B) B220^+^NP^+^ B cells. (C) Frequency of B220^+^GL-7^+^CD95^+^ GC B cells of total B220^+^ B cells. (D) Frequency of B220^+^GL-7^+^CD95^+^NP^+^ GC B cells of total B220^+^ B cells. (E) Representative flow plot of total IgD^−^CD138^+^TACI^+^ plasma cells. (F) Gating strategy for different subsets of PCs. (G) Percentages of subsets of plasma cells (total IgD^−^CD138^+^TACI^+^ plasma cells, IgD^−^CD138^+^TACI^+^B220^int^CD19^+^ plasmablasts, IgD^−^CD138^+^TACI^+^B220^−^CD19^+^ immature plasma cells, IgD^−^CD138^+^TACI^+^B220^−^CD19^−^ mature/resting plasma cells). (H) Representative flow and dot plots of NP^+^IgD^−^CD138^+^TACI^+^ plasma cells. Frequencies of (I) CD4^+^CD44^+^CD62L^−^ splenic effector and (J) CD4^+^CD44^+^CD62L^−^CXCR5^+^PD-1^+^ Tfh of total CD4^+^ T cells. (K) High affinity (NP4) and (L) low affinity (NP28) anti-NP and total serum IgG and IgG1 titers. Each symbol represents an individual mouse (n =7-10 mice per group) and data are presented as means ± SEM. Data in each panel represent two experiments. P values were calculated via an unpaired Student’s t-test (A-J) or two-way Anova with Dunn-Sidak correction (K-L) (NS, not significant, p>0.05, *, p <0.05, **, p <0.01).

Together, findings from SLE-associated and immunization-induced responses highlight a critical role of IRF7 in promoting systemic autoimmune responses but not foreign antigen-induced GC, PC and antibody responses.

### IRF7 expression in the hematopoietic compartments drives spontaneous GC and PC responses in FcγRIIB^−/−^ mice

To determine the effects of IRF7 expression in the host microenvironment versus hematopoietic compartment on autoimmune GC and PC responses in FcγRIIB^−/−^ mice, we generated competitive mixed bone marrow chimeras. Lethally irradiated wild-type hosts (CD45.1^+^) received a 50:50 mix of bone marrow from FcγRIIB^−/−^ (CD45.1^+^CD45.2^+^) and FcγRIIB^−/−^ IRF7^−/−^ (CD45.2^+^) donors (Figure 4A). Eight weeks post-reconstitution recipient mice were equally reconstituted by both donor populations (Figure 4B). Analysis of B cell populations revealed no significant differences in the development of naïve follicular B cells (Figure 4C, D) or ABCs (Figure 4E, F). However, we observed a significant reduction in IRF7-deficient B cells within splenic GC (Figure 4G, H), and PC responses within spleen and bone marrow (Figure 4I, J). Interestingly, IRF7 deficiency in hematopoietic cells did not affect Teff and Tfh populations (Figure 4K, L), and numerous myeloid populations except for iDCs which remained significantly reduced and pDCs which were elevated in the absence of IRF7 (Figure 4M). These findings suggest the importance of IRF7 expression in hematopoietic cells in B cell differentiation into GC B cells and PCs during systemic autoimmune responses in FcγRIIB^−/−^ mice.

**Figure 4:**
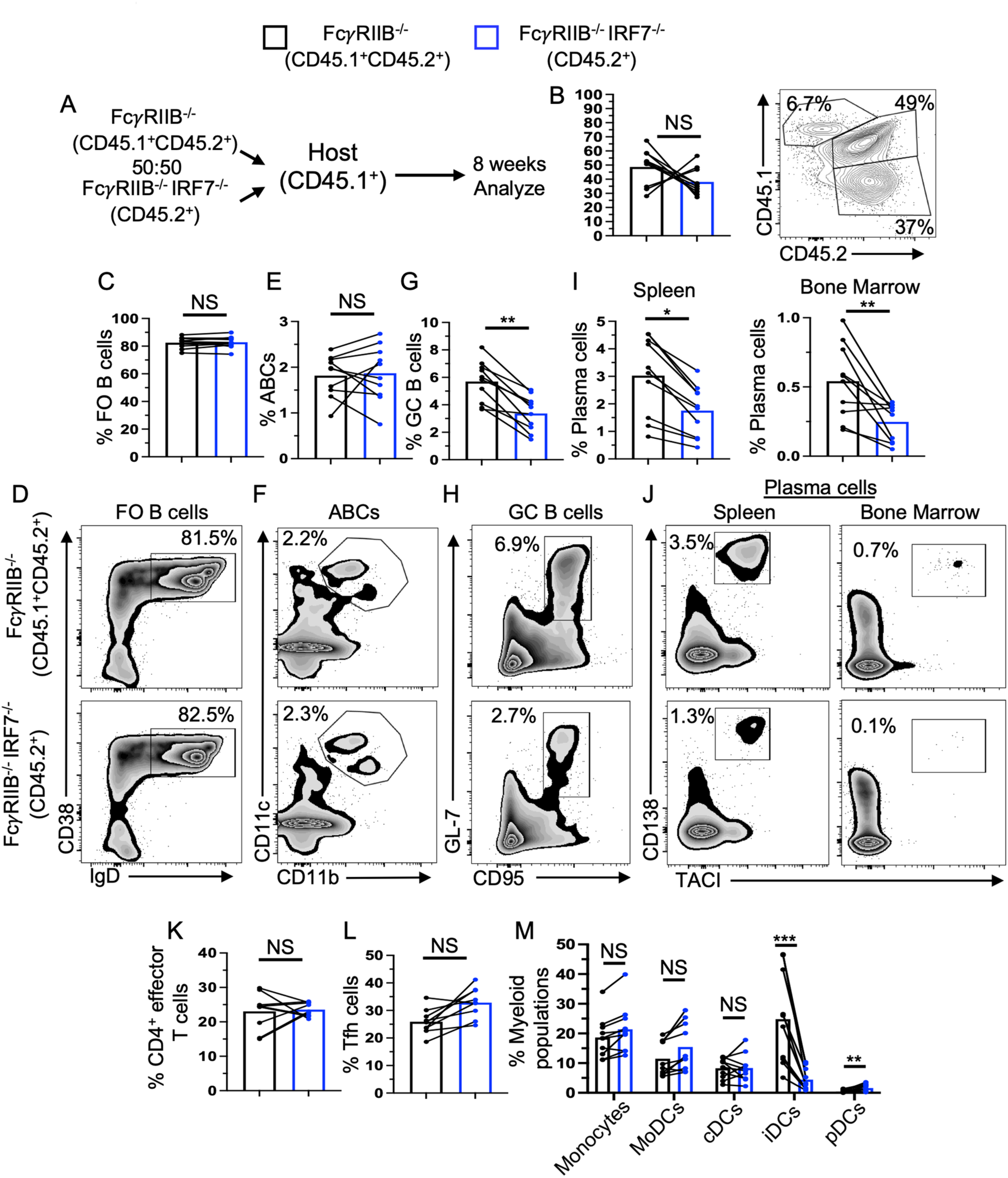
IRF7 controls autoimmune GC and AFC responses by functioning in hematopoietic cells. (A) The schematic of competition chimeras in which allotype marked bone marrow from FcγRIIB^−/−^ (CD45.1^+^CD45.2^+^) and FcγRIIB^−/−^IRF7^−/−^ (CD45.2^+^) mice were transferred into lethally irradiated CD45.1^+^ hosts. (B) The bars and flow cytometry plot show bone marrow reconstitution of FcγRIIB^−/−^ (CD45.1^+^CD45.2^+^) and FcγRIIB^−/−^IRF7^−/−^ (CD45.2^+^) donor cells. Flow cytometry analysis of (C) the frequency and (D) gating strategy of CD38^+^IgD^+^ FO B cells of total B220^+^CD19^+^ splenic B cells. Flow cytometry analysis of (E) the frequency and (F) gating strategy of IgD^−^CD11b^+^CD11c^+^ ABCs of B220^+^CD19^+^ total splenic B cells and (G) the frequency and (H) gating strategy of B220^+^GL-7^+^CD95^+^ GC B cells of B220^+^CD19^+^ total splenic B cells. Flow cytometry analysis of (I) the frequency and (J) gating strategy of CD138^+^TACI^+^ splenic and bone marrow plasma cells of IgD^−^ B cells (shown in Figure 1H). Flow cytometry analysis of the frequencies of splenic (K) CD4^+^ effector T cells, and (L) Tfh of total CD4^+^ cells and (M) the frequencies of various myeloid cells of total myeloid populations. Each symbol represents an individual mouse (n = 10-14 mice per group) and data are presented as means ± SEM. Data in each panel represent three experiments. P values were calculated via an unpaired Student’s t-test (Not significant, NS, p >0.05, *, p <0.05, **, p <0.01, ***, p <0.001).

### Single cell transcriptome of SLE-prone B cells identifies 14 clusters and their regulation by IRF7-driven transcriptional programs

Given the effects of IRF7 expression in hematopoietic cells on spontaneous B cell responses, we next examined the role of IRF7 in transcriptional control of B cell responses in SLE by performing scRNAseq on splenic B cells sorted from 3 month old FcγRIIB^−/−^IRF7^−/−^ and FcγRIIB^−/−^ mice. To be able to detect transcriptome in subsets of splenic activated B cells that have small percentage representations compared to naïve B cells, analysis was performed on B cells composed of 90% IgD^−^ antigen-experienced B cells and 10% IgD^+^ naïve B cells (Figures 5A, S5A). Unsupervised clustering of 36,842 B cells passing QC identified 14 B cell clusters as visualized using UMAP (Figures 5B, S5B, C). These B cell clusters were annotated based on the expression of canonical markers specific for subpopulations of B cells (Figure 5C) and by comparing our new data set with the annotated scRNAseq previously reported by others (Laidlaw et al., 2020). We identified a sharp reduction in the percentage of IRF7^−/−^ B cells in the pre-GC transition, GC dark (DZ) and light (LZ) zone and plasma cell/blast clusters, but not in the follicular cluster 2, marginal zone, or switched activated populations (Figure 5D, E). Transitional, follicular cluster 1 and naïve activated B cells were proportionally higher in IRF7^−/−^ B cells than control cells (Figure 5E). In summary, scRNAseq of autoimmune B cells sufficient and deficient in IRF7 identified 14 clusters and revealed significant differences in subpopulations of B cells, pointing to potential regulation of autoimmune GC and PC responses by IRF7-driven transcriptional programs.

**Figure 5:**
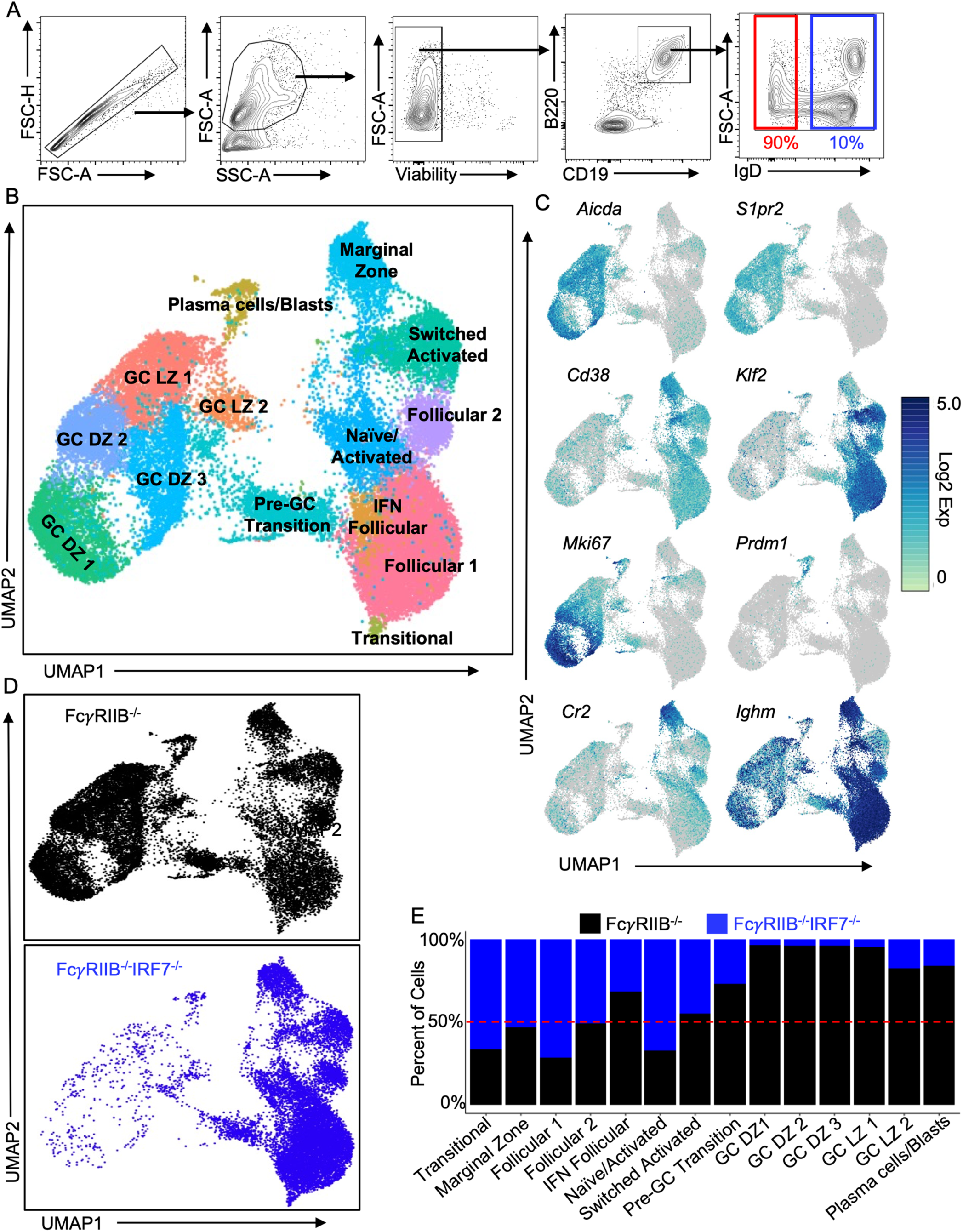
Single cell RNAseq identifies 14 clusters in SLE-prone FcγRIIB^−/−^ B cells. (A) Gating strategy for sorting 90% IgD^−^ activated and 10% IgD^+^ naïve B cells for scRNAseq analysis. (B) Unsupervised clustering of 36,842 splenic B cells from 3 mo old FcγRIIB^−/−^ and FcγRIIB^−/−^IRF7^−/−^ mice, visualized using UMAP. Cells analyzed contained 10% B220^+^IgD^+^ naïve and 90% B220^+^IgD^−^ activated B cells that were sorted from two individual FcγRIIB^−/−^ and FcγRIIB^−/−^IRF7^−/−^ mice (n = 4 mice). (C) Representative UMAPs of hallmark genes to demonstrate annotation of populations in the scRNAseq dataset. Gene expression for clusters is shown at a log scale. (D) UMAPs depicting the distribution of cells between the FcγRIIB^−/−^ and FcγRIIB^−/−^ IRF7^−/−^ mice. (E) The percentage of cells in each B cell cluster from FcγRIIB^−/−^ and FcγRIIB^−/−^IRF7^−/−^ mice is shown in the bar graphs.

### IRF7-driven transcriptome in B cells regulates B cell differentiation through pre-GC, GC and PC populations during a systemic autoimmune response

To determine the role of IRF7 in B cell differentiation into the GC and PC pathways, we analyzed the top 100 genes differentially expressed between IRF7 sufficient and deficient B cells across the scRNAseq dataset and identified dynamic transcriptional changes associated with B cell differentiation (Figure 6A). To further investigate the involvement of IRF7 in regulating B cell differentiation, we used pseudotime inference to identify transcriptional programs associated with B cell differentiation (Figure 6B). The pseudotime trajectory revealed that B cells differentiated in two major pathways: one group differentiated through pre-GC transitional bridge into GC and PC clusters, and the other group into naïve activated and switched activated B cells and marginal zone B cells (Figure 6B) which can contribute to the generation of autoantibody producing cells and autoantibodies. We found that IRF7 deficiency in B cells predominantly affected B cell differentiation through pre-GC cluster into GC and PC clusters. We further re-clustered the pre-GC transitional B cell cluster from our scRNAseq dataset to identify underlying subpopulations and their regulation by IRF7 (Figure 6C). The re-clustered pre-GC transitional B cells produced 5 subclusters that allowed further examination. We examined these 5 subclusters for gene markers associated with B cell differentiation and manually annotated the 5 subclusters based on cluster-based gene expression. Pre-GC1 expressed numerous genes that distinguished them from naïve follicular B cells, but they retained most follicular markers in the subcluster. As B cells differentiated through pre-GC stages, they downregulated follicular B cell markers (*Cd38*, *Klf2*, *S1pr1*) and began to upregulate markers of activation/GC differentiation (Figure 6D, *S1pr2*, *Aicda*, *Mef2b*, *Mef2c*, *Pou2f2*, *Ezh2*, *Fas*, *J chain*, and *Slamf7*). Further analysis of scRNAseq data indicated that cells of the pre-GC1 cluster were quiescent followed by proliferation of B cells at pre-GC2 and pre-GC3 clusters (Figure 6E, G). Finally, we determined the percentage of each subcluster that was composed of FcγRIIB^−/−^ or FcγRIIB^−/−^IRF7^−/−^ B cells and found a stark reduction in FcγRIIB^−/−^IRF7^−/−^ B cells starting at the pre-GC2 cluster (Figure 6E, F). These data indicate that IRF7-driven genetic programs regulate B cell differentiation through pre-GC, GC, and PC fates in SLE-prone FcγRIIB^−/−^ mice.

**Figure 6:**
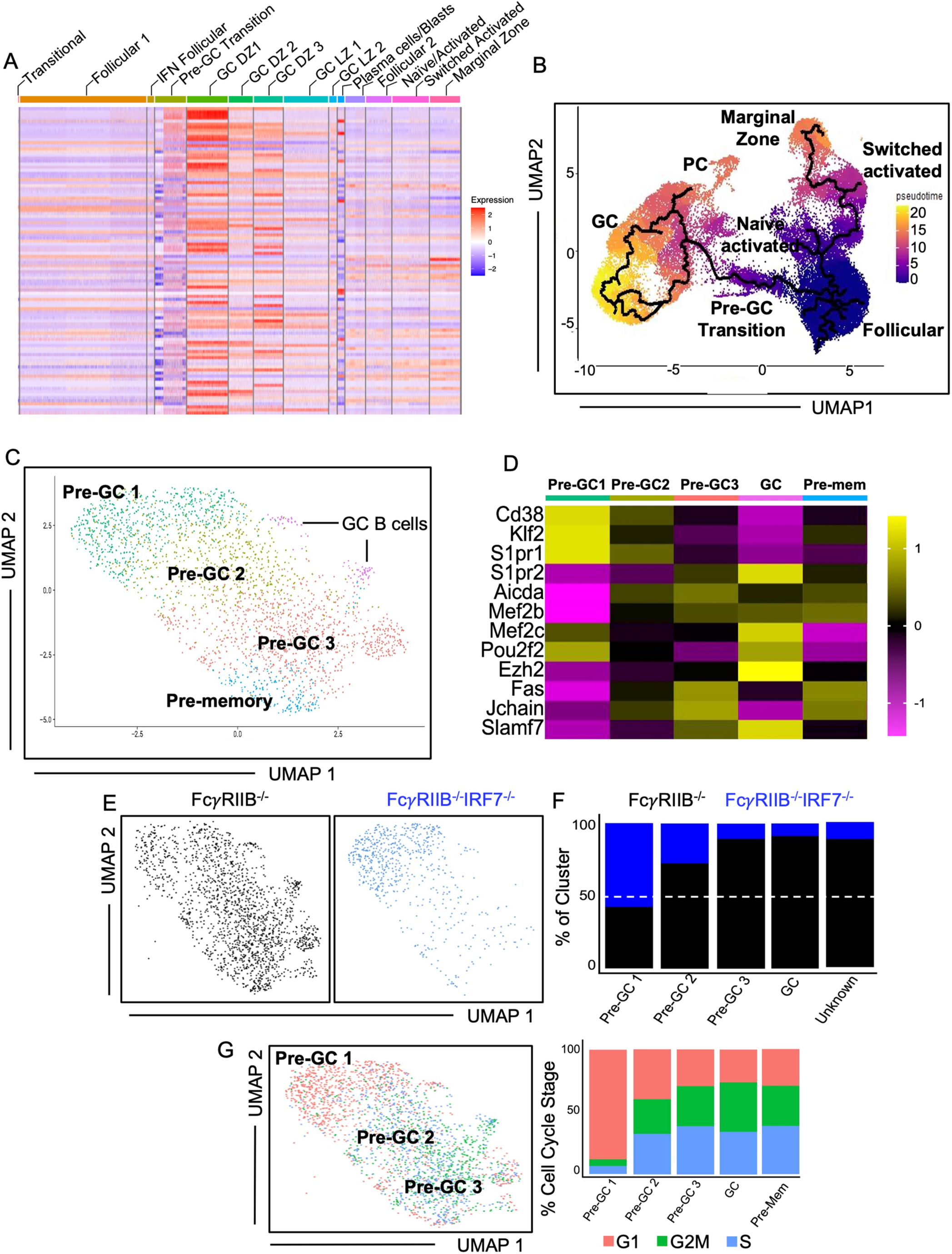
Single cell RNAseq delineates the role of IRF7 in B cell differentiation through GC and plasma cell fates. (A) Top 100 differentially expressed genes across the scRNAseq dataset were analyzed among all clusters. (B) Pseudotime analysis of the scRNAseq dataset projected using the UMAP to transcriptionally predict the differentiation projection of follicular B cells in FcγRIIB^−/−^ mice. (C) Subclustering of the pre-GC transitional cluster (n = 2,461 cells) of splenic B cells from FcγRIIB^−/−^ mice using (D) markers associated with B cell differentiation into GC B cells. (E) Cell cycle analysis of cells in these subclusters. (F) The cells in subclusters were compared between FcγRIIB^−/−^ and FcγRIIB^−/−^IRF7^−/−^ mice. (G) The percentages of cells in each subcluster from FcγRIIB^−/−^ and FcγRIIB^−/−^IRF7^−/−^ mice that are in cell cycle are shown in the bar graphs.

### IRF7 promotes systemic autoimmunity by regulating transcription, translation and metabolism in activated B cells

To understand the B cell-intrinsic mechanism of IRF7-mediated systemic autoimmunity, we performed pathway analysis on differentially expressed genes identified between FcγRIIB^−/−^ and FcγRIIB^−/−^IRF7^−/−^ mice within the pre-GC clusters of the scRNAseq. The top two pathways identified contained translation (EIF2 signaling) and Oxidative Phosphorylation (OXPHOS) related genes including ribosomal genes that were differentially expressed between FcγRIIB^−/−^ and FcγRIIB^−/−^IRF7^−/−^ pre-GC clusters (Figure 7A, B). We also performed IRF7 ChIP-seq on antigen-experienced IgD^−^ B cells from FcγRIIB^−/−^ mice and identified 1229 IRF7 peaks and approximately 32% mapped within 3kb of a promoter and the remainder fell largely within intergenic regions or introns (Figure 7C). Combined scRNAseq and ChIPseq analysis identified overlap between 42 upregulated and 29 downregulated genes between FcγRIIB^−/−^ and FcγRIIB^−/−^ IRF7^−/−^ B cells that were IRF7 targets (Figure 7D). Surprisingly, we found IRF7 peaks in the promoters of several ribosomal protein genes (Figure 7E). To validate the OXPHOS pathway identified in the scRNAseq data set (Figure 7A), we performed extracellular flux analysis which revealed lower OXPHOS and glycolysis in FcγRIIB^−/−^IRF7^−/−^ B cells compared to FcγRIIB^−/−^ control B cells including basal respiration and non-mitochondrial O2 consumption rate (Figure 7F, G). To validate the EIF2 signaling pathway (Figure 7A) linked to translation, we measured translation and found that FcγRIIB^−/−^IRF7^−/−^ GC B cells and PCs had reduced translation compared to FcγRIIB^−/−^ control B cell counterparts (Figure 7H). FcγRIIB^−/−^IRF7^−/−^ B cells also showed reduced translation compared to FcγRIIB^−/−^ control B cells post-TLR7 stimulation but not IgM and CD40 (Figure 7I). These data suggest that IRF7 transcriptionally regulates translation and metabolism in B cells, promoting SLE-associated GC and PC responses.

**Figure 7:**
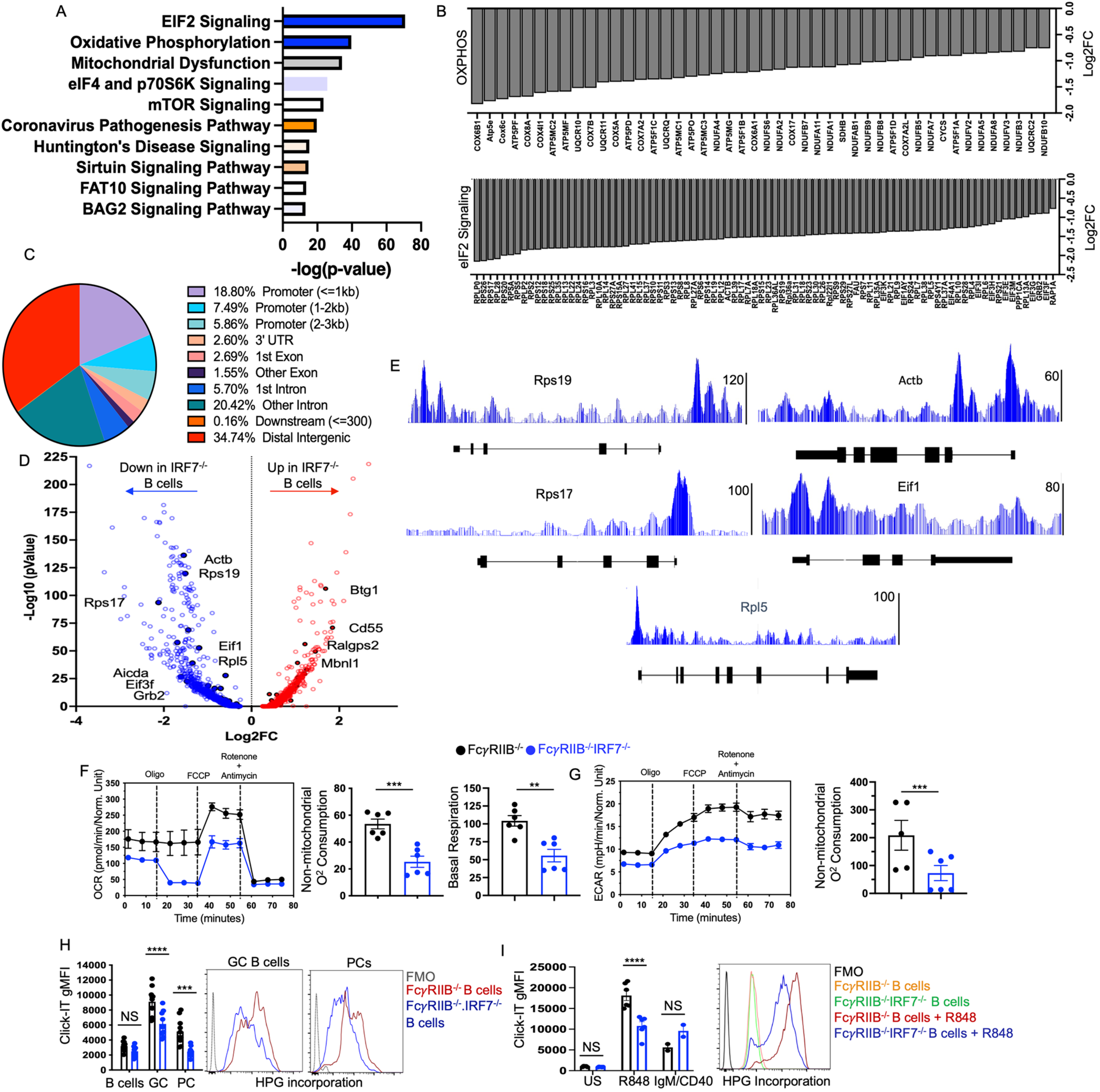
IRF7 regulates translation and metabolism during B cell activation. **(A)** Ingenuity pathway analysis of differentially expressed genes within the pre-GC clusters of the scRNAseq data from FcγRIIB^−/−^ and FcγRIIB^−/−^IRF7^−/−^ mice. (B) The heatmaps of upregulated and downregulated genes linked to EIF2 signaling and Oxidative Phosphorylation (OxPhos) between FcγRIIB^−/−^ and FcγRIIB^−/−^IRF7^−/−^ pre-GC clusters. (C) The pie chart showing the genomic locations of IRF7 peaks from ChIP-seq analysis of antigen-experienced IgD^−^ B cells from FcγRIIB^−/−^ mice. (D) The volcano plot of IRF7 target genes identified by ChIPseq analysis which are upregulated and downregulated between FcγRIIB^−/−^ and FcγRIIB^−/−^IRF7^−/−^ B cells in scRNAseq analysis. (E) Representative IRF7 peaks in the promoters of several ribosomal protein genes. (F, G) Extracellular flux analysis showing OXPHOS and glycolysis of FcyRIIB^−/−^IRF7^−/−^ and FcyRIIB^−/−^ B cells including basal respiration and non-mitochondrial O2 consumption rate. (H) Translation analysis in total and spontaneously activated B cells from FcγRIIB^−/−^ and FcγRIIB^−/−^ IRF7^−/−^ mice. (I) Analysis of translation in TLR7-activated B cells from FcγRIIB^−/−^ and FcγRIIB^−/−^ IRF7^−/−^ mice by flow cytometry. Data in (A-E) represent 4 mice. Each symbol in (F-I) represents an individual mouse (n=3-6 mice group) and data are presented as means ± SEM. Data in (F-I) represent two-four experiments. P values were calculated via an unpaired Student’s t-test (F-G) or two-way Anova with Dunn-Sidak correction (H, I) (Not significant, ns, p <0.05, **, p <0.01, ***, p <0.001, ****, p <0.0001).

### B cell-intrinsic IRF7 is required for loss of tolerance and autoantibody production but not sufficient for promoting overall spontaneous GC and PC responses

Given the effects of IRF7 on B cell transcriptome, translation and metabolism, we measured IRF7 levels in B cell subsets in FcγRIIB^−/−^ mice. IRF7 expression was elevated in pre-GC and GC B cells relative to non-GC B cells with the highest expression in GC B cells and much higher expression in FcγRIIB^−/−^ B cells than B6 control B cells (Figure 8A, B). We also analyzed the published RNAseq dataset (Jenks et al., 2018) and found higher IRF7 expression in SLE B cells than healthy control B cells (Figure 8C). In addition, we analyzed IRF7 expression in transitional 3 (T3), resting naïve (rN), activated naïve (aN), double negative (DN2) B cells, switched memory (SM), and plasmablast/plasma cells (ASCs) from healthy and SLE samples. *IRF7* expression was increased in activated B cells in SLE patients (Figure 8D) and during activation of B cells from healthy PBMCs for 24 hours with a TLR7 agonist (Figure 8E).

**Figure 8:**
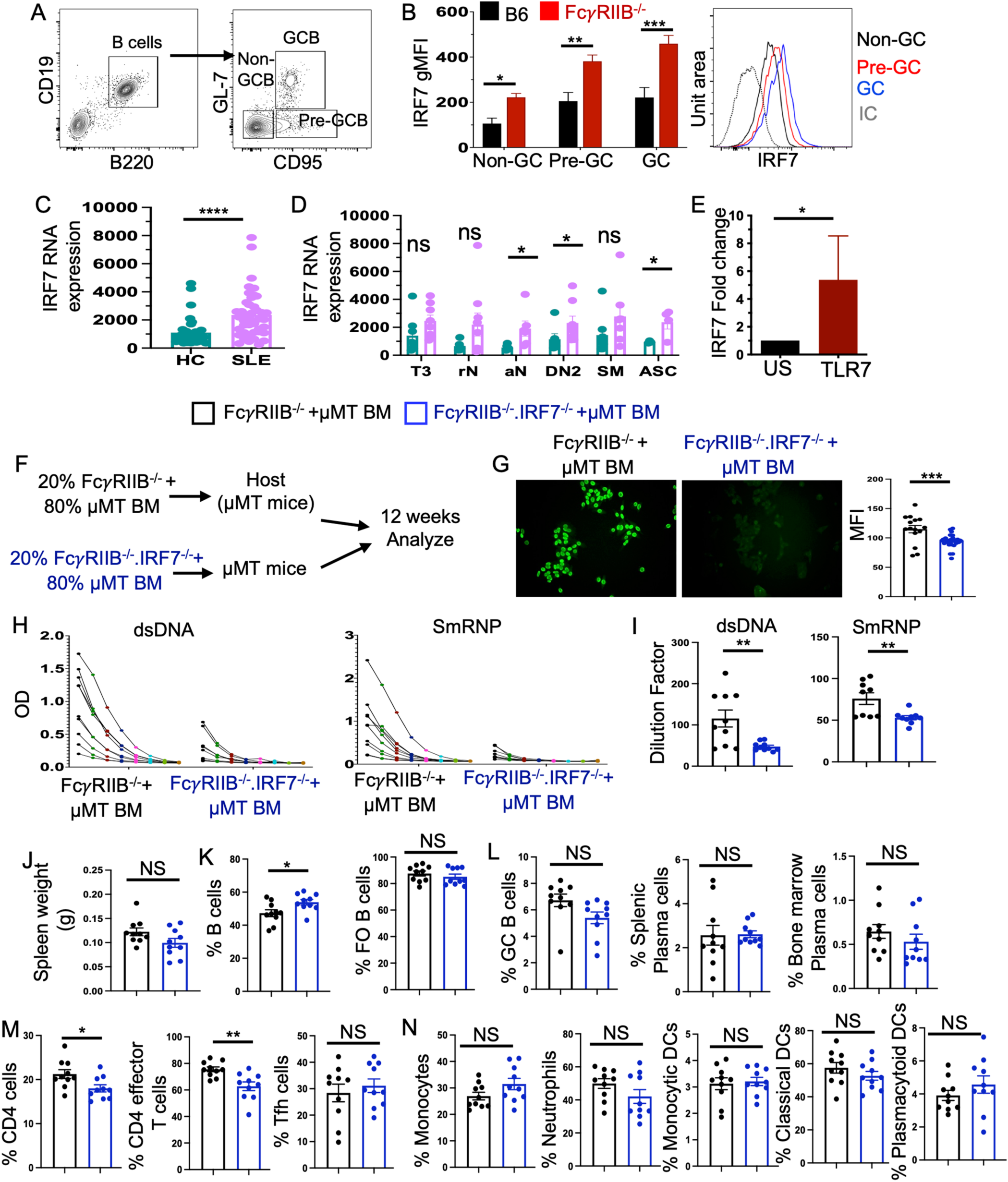
B cell-intrinsic and -extrinsic roles of IRF7 in breaking tolerance and autoimmune PC and GC responses. (A) Gating strategy of B220^+^CD19^+^GL-7^−^CD95^−^ non-GC B cells (non-GCB), B220^+^CD19^+^GL-7^−^CD95^+^ pre-GC B cells (pre-GCB) and B220^+^CD19^+^GL-7^+^CD95^+^ GC B cells (GCB). (B) IRF7 expression in gMFI in non-GCB, pre-GCB and GCB cells in B6 and FcγRIIB^−/−^ mice. Our analysis of published RNAseq data of healthy control (HC) and systemic lupus erythematosus (SLE) patients showing IRF7 RNA expression in total (C) and various subsets of B cells (D) such as transitional 3 (T3), resting naïve (rN), activated naïve (aN), double negative (DN2) B cells, switched memory (SM), and plasmablast/plasma cells (ASCs). (E) *IRF7* expression was measured in B cells from healthy PBMCs following activation for 24 hours with a TLR7 agonist. (F) Schematic of B cell-specific mixed bone marrow chimeric mice generation. (G) Immunofluorescent staining of Hep-2 slides for serum ANA seropositivity and (H, I) serum anti-dsDNA and anti-SmRNP antibody titers in μMT recipient mice with FcγRIIB^−/−^ B cells sufficient and deficient IRF7. RNA expression in healthy control (HC) and SLE patient B cells. (J-L) Spleen weight and frequencies of B220^+^ total and FO B cells, GC B cells, splenic plasma and bone plasma cells of B220^+^ total B cells. (M) Flow cytometry analysis of frequencies of CD4^+^ total T cells, CD4^+^ effector T cells and Tfh cells of total CD4^+^ T cells. (N) Flow cytometry analysis of frequencies of splenic myeloid cells in these mice. Each symbol represents an individual mouse (n = 9-10 mice per group) and data are presented as means ± SEM. Data in each panel represent two independent experiments. P values were calculated via two-way Anova with Dunn-Sidak correction (B) or an unpaired Student’s t-test (C-E and G-N) (NS, not significant, *, p <0.05, **, p <0.01, ***, p <0.001, ****, p <0.0001).

Next to determine whether IRF7 functioned in a B cell-intrinsic or -extrinsic manner to promote spontaneous autoimmune GC and PC responses and autoantibody production, we generated B cell-specific mixed bone marrow (BM) chimeras. BM cells from FcγRIIB^−/−^ or FcγRIIB^−/−^IRF7^−/−^ mice were mixed with BM cells from μMT mice, that lack B cells, in a 20:80 ratio and transferred into lethally irradiated μMT recipient mice as depicted in Figure 8F. Serum ANA positivity and dsDNA- and SmRNP-reactive autoantibody titers in recipient mice with B cells deficient in IRF7 were significantly reduced compared to recipient mice with B cells sufficient in IRF7 (Figure 8G-I). However, no major differences in spleen weight (Figure 8J) and frequencies of naïve (Figure 8K) and activated (Figure 8L) B cell populations were observed between the strains except for an increased percentage of total B cells in the absence of IRF7 in B cells. B cell-specific IRF7-deficient mice had reduced frequencies of total CD4 and CD4 effector T cells, although no differences in the frequency of Tfh (Figure 8M) and myeloid cells (Figure 8N) were observed between the strains. These data indicate a B cell-intrinsic requirement of IRF7 in loss of immune tolerance, autoantibody production to DNA and RNA associated autoantigens and CD4^+^ effector T cell responses whereas both B cell-intrinsic and -extrinsic contributions of IRF7 are required for driving GC, PC and Tfh cell responses in SLE-like autoimmunity.

## Discussion

Despite a longstanding association of IRF7 with SLE, the cell-intrinsic role and mechanism by which IRF7 promotes spontaneous SLE-associated GC and PC responses that harbor autoreactive B cells in SLE are not known. Here using IRF7-deficient mice, competitive mixed bone marrow (BM) and B cell-specific BM chimeras, we identified previously unknown B cell-intrinsic and -extrinsic roles for IRF7 in promoting spontaneous autoimmune GC and PC responses. We found that IRF7 is involved in B cell differentiation into autoantibody producing cells, steady-state total and ANA-specific class-switched IgG antibody responses and SLE pathogenesis. IRF7, however, was dispensable for promoting foreign antigen driven GC, PC and antibody responses. Through the single cell transcriptome analysis of SLE-prone B cells, we uncovered the functions for IRF7-driven transcriptional programs in B cell differentiation through pre-GC into GC and PC pathways during systemic autoimmune responses. Analysis of ChIPseq data identified IRF7 target genes that are differentially expressed in IRF7-deficient SLE-prone B cells. Mechanistically, IRF7 promotes systemic autoimmunity by regulating B cell transcriptome, translation, and metabolism required for B cell activation and differentiation into GC and PC populations during SLE-like autoimmune responses.

Previous studies highlighted the roles of both the extrafollicular (EF) (Jenks et al., 2018; William et al., 2002) and GC pathways (Arkatkar et al., 2017; Cappione et al., 2005; Degn et al., 2017; Jackson et al., 2016; Soni et al., 2014; Tiller et al., 2010; Vinuesa et al., 2009) in the development of autoreactive B cells and pathogenic Ab production in SLE-prone mice and patients. In particular, using the FcγRIIB^−/−^ SLE mouse model used in the current study, Tiller et al. previously showed the development of autoreactive GC B cells and PCs in which IgG autoantibodies were enriched in GCs and somatically hypermutated (Tiller et al., 2010), highlighting the role of the GC pathway in pathogenic autoantibody production. By blocking the GC pathway previous studies highlighted a pathogenic role for the EF pathway in systemic autoimmunity (Brown et al., 2022b; Voss et al., 2022). We observed dampened GC and PC responses, low numbers of autoantibody-producing splenic and bone marrow AFCs and reduced serum ANA titers in FcγRIIB^−/−^ mice in the absence of IRF7. Together, our current findings and previously described role of the GC pathway in pathogenic Ab production in FcγRIIB^−/−^ mice (Tiller et al., 2010) strongly suggest the role of IRF7 in the generation of autoreactive PCs and pathogenic Abs through the GC pathway. Several studies have demonstrated that ABCs generated through EF pathways, can either differentiate into EF plasmablasts or participate within the GC (Nickerson et al., 2023; Ricker et al., 2021). Deeper understanding of the differential mechanisms by which GC and EF pathways promote high affinity autoantibody-producing PCs in SLE is warranted to develop targeted therapeutics.

Previous studies have delineated the contributions of IRF5, including a B cell-intrinsic role of IRF5, in promoting systemic autoimmunity and disease (Ban et al., 2021; Banga et al., 2020; Cham et al., 2012; Pellerin et al., 2021; Richez et al., 2010). Here, for the first time, we identified the role of IRF7 in promoting PC and GC response and autoantibody production. It is thought that there is a significant overlap between IRF5 and IRF7 functions; however, it is not clear whether IRF7 homodimerizes or heterodimerizes with IRF5 or other IRF family members (i.e., IRF3) in B cells to promote GC and PC responses in systemic autoimmunity. The family of IRF proteins is well-known to participate in transcriptional coregulation of gene expression (Andrilenas et al., 2018). Studies focusing on the dynamic interactions and the transcriptional landscape regulated by IRF proteins singularly or cooperatively will help develop a deeper understanding of the mechanisms of IRF7 mediated regulation of B cell responses in SLE.

Through the generation of B cell-specific BM chimeras, we show here a B cell-intrinsic requirement of IRF7 in loss of tolerance and autoantibody production. Our data suggest that B cell-intrinsic IRF7 controls GC and/or PC tolerance checkpoints of autoreactive B cell selection. Our bone marrow chimeric data, OMICS and mechanistic data together suggest that IRF7-regulated transcriptional programs, translation and metabolic reprogramming in B cells play critical roles in breaking tolerance and autoreactive B cell selection. B cell-intrinsic IRF7 deficiency, however, was not sufficient to affect overall spontaneous PC, GC and Tfh responses. These data suggest that development of PC, GC and Tfh responses in the BM chimeric environment requires both B cell-intrinsic and -extrinsic IRF7. Although correlative, IRF7 is thought to contribute to SLE in a B cell-extrinsic manner by inducing T1-IFNs in pDCs (Honda et al., 2005; Swiecki and Colonna, 2015). Whether B cell-extrinsic IRF7 contributions to autoimmune PC, GC and Tfh responses in SLE models are mediated through pDC-derived T1- IFN signaling will require an in vivo system, not yet readily available, in which IRF7 can be deleted in pDCs.

Several risk variants in and around IRF7 are associated with SLE disease activity (Fu et al., 2011; Salloum et al., 2010). A mutation in the IRF7 gene resulting in Q412R is associated with elevated IRF7 expression and an increased T1-IFN signature in SLE patients (Fu et al., 2011). However, it has remained unclear whether the IRF7 risk allele contributed to autoimmune GC and PC responses in SLE in a cell-intrinsic manner independent of T1-IFN production. Generation of mouse models carrying IRF7 risk variants will help address this unanswered question. Here we have demonstrated that outside of the T1-IFN genes, IRF7 can regulate genes in GC B cells and PCs that are involved in translation and metabolism, which are critical for B cell differentiation through GC and PC pathways in a systemic autoimmune response.

A previous study demostrated the role of IRF7 in promoting autoantibody responses in a chemically (pristane)-induced B6 model of SLE (Miyagawa et al., 2016). Surprisingly, despite reductions in IC deposition in the kidney of IRF7^−/−^ mice, kidney pathology, and proteinuria were reported in these mice (Miyagawa et al., 2016). The role of IRF7 in autoimmune GC and PC responses and disease in spontaneous SLE models, which may involve differential mechanism to drive autoimmunity, has remained unclear. We observed reductions in autoantibody production, IC deposition, kidney pathology and serum BUN/Creatinine levels in IRF7 deficient spontaneous SLE-prone mice. We also observed significantly improved survival in TLR7-accelerated SLE-prone mice deficient in IRF7. Our data suggest that IRF7 plays a major role in promoting GC and PC responses and systemic autoimmunity including tissue inflammation and disease. The discrepancy between our findings and the previous study could be due to differences in the involvement of differential mechanisms between the spontaneous SLE-prone versus chemically induced non-autoimmune B6 mouse models that were used.

We utilized scRNAseq analysis to profile subpopulations of B cells and their differentiation trajectories in SLE-prone mice. Pseudotime trajectory analysis of differentiating B cells revealed that during a systemic autoimmune response, B cells differentiate into two distinct pathways in which one group differentiated through pre-GC transitional bridge into GC and PC fates, and the other group into naïve activated and switched activated B cells. By comparing single-cell transcriptomic data of FcγRIIB^−/−^ and FcγRIIB^−/−^IRF7^−/−^ B cells, we discovered a significant deficit in IRF7^−/−^ B cell differentiation into GC and plasmblasts/PC clusters. Through targeted re-clustering of pre-GC transitional cells, we further discovered that B cell differentiation was halted at the pre-GC1 stage in the absence of IRF7 in which majority of these cells were quiescent compared to B cells in FcγRIIB^−/−^ mice that progressed to pre-GC2 and pre-GC3 stages were in cell cycle. Our OMICs and validation data together suggest that during autoimmune B cell responses in spontaneous SLE-prone mice, IRF7 induces transcription of genes required for protein translation and metabolism in B cells. These IRF7-driven events in B cells together with B cell-extrinsic factors, likely T1-IFN signaling, are required to meet the demands of activated B cells that can differentiate into GC and PC fates, promoting the generation of pathogenic autoantibody-producing cells and leading SLE development.

### Limitations of the study

Our identification of critical B cell-intrinsic and -extrinsic roles for IRF7 in regulating their differentiation into the GC and PC pathways does not negate the potential IRF7 role in regulating other B cell populations. Although we focused on the role of IRF7 in B cell differentiation into the GC/PC clusters, we identified differentially expressed genes in the naïve activated, switched activated and marginal zone (MZ) B cell clusters that will be a topic of further investigation at the mechanistic level. Additionally, based on our current findings suggesting both B cell-intrinsic and -extrinsic roles for IRF7 in autoimmune GC and PC responses and in loss of B cell tolerance in systemic autoimmunity, we primarily focused our investigations into the IRF7 role in B cell responses. We, however, cannot exclude the possibility that IRF7 may contribute to the functionality of myeloid or T cell populations and thereby contributing to the autoimmune B cell response and autoimmunity. Thus, interrogation of IRF7 functions in other cell types in promoting autoimmune GC and PC responses and systemic autoimmunity also will be necessary in the future.

## MATERIALS AND METHODS

### Study Design

The overall goal of this study was to delineate the cell-intrinsic and -extrinsic mechanisms by which IRF7 promotes autoimmune B cell responses in systemic autoimmunity. This was accomplished by first generating IRF7-deficient SLE-prone mice and then by generating competitive and B cell-specific mixed BM chimeras. The phenotypic examination of various B and T cell populations were performed by flow cytometry and immunofluorescence microscopy. Mechanistic studies were performed through single cell transcriptome and ChIPseq of B cells from SLE-prone mice sufficient and deficient in IRF7. The number of mice per experiment was determined based on the power calculation of our published data (Chodisetti et al., 2020; Domeier et al., 2018; Fike et al., 2021). Littermate controls were used where possible. Control and experimental mice were age- and sex-matched. All data points generated in various experiments were included in the analysis. All experimental findings were replicated, and the number of replicates is indicated in the figure legends.

### Mice

C57Bl/6 (B6), B6.SJL-*Ptprc^a^Pepc^b^/BoyJ* (CD45.1) and B6.129S2-Ighm^tm1Cgn^/J (B6.μMT), mice were originally purchased from the Jackson laboratories and bred in house. B6.129P2- *Irf7*^tm1Ttg^/TtgRbRc (IRF7^−/−^) mice were purchased from Riken BRC and bred internally. FcγRIIB^−/−^ yaa mice were generated by crossing FcγRIIB^−/−^ mice described previously (Bolland et al., 2002; Shinde et al., 2018) to B6.yaa mice (from the Jackson Laboratories) internally. All crosses generated for this manuscript were performed in house. All animal studies were conducted at Pennsylvania State University Hershey Medical Center in accordance with the guidelines approved by our Institutional Animal Care and Use Committee. Animals were housed in a specific pathogen-free barrier facility.

### Generation of mixed bone marrow chimeras

#### Mixed competition bone marrow (BM) chimeric mice

These mice were generated by lethally irradiating 10–12-week-old female CD45.1 hosts in two doses of 450 rads (X-RAD 320iX Research Irradiator; Precision X-Ray) within a 4h interval. Within a few hours of irradiation, each recipient mouse received 10×10^6^ T cell-depleted BM cells intravenously composed of 5×10^6^ cells from FcγRIIB^−/−^ CD45.1^+^/CD45.2^+^ and 5×10^6^ cells from FcγRIIB^−/−^IRF7^−/−^CD45.2^+^ donors (50:50). Recipient mice were analyzed 8 weeks post-BM reconstitution.

#### B cell-specific mixed BM chimeras

8–10-wk-old female B6.μMT (B6.129S2-Igh^mtm1^Cgn/J) recipient mice were lethally irradiated with 900 rads of x rays (X-RAD 320iX Research Irradiator; Precision X-Ray). Within 4-6h post-irradiation, each recipient intravenously received 10^7^ T cell–depleted BM cells isolated from 8–10-wk-old female donor mice with 80% of cells from B6.μMT mice and 20% from FcγRIIB^−/−^ or FcγRIIB^−/−^IRF7^−/−^ mice. Recipients were analyzed after 3 months.

#### Immunization of mice

Eight to ten wk old age and sex matched mice were immunized intraperitoneally (i.p.) with 200µg of 4-Hydroxy-3-nitrophenylacetyl-Keyhole Limpet Hemocyanin (NP-KLH, Biosearch Technologies) mixed with Complete Freund’s Adjuvant (CFA, Sigma-Aldrich). Spleens were harvested and processed for various analysis on indicated day post immunization.

### Flow cytometry

Analysis of splenic populations was performed by processing the tissue into a single cell suspension followed by RBC lysis using Tris-Ammonium Chloride. Splenocytes were stained using the antibodies listed in the table below. For bone marrow analysis, cells were flushed from isolated femurs with RPMI followed by RBC lysis and staining. For the flow cytometry of kidney, the kidneys were perfused prior to harvesting. Kidneys were minced into small pieces using a scalpel and incubated with collagenase Type I (Sigma) for 30 mins at 37°C prior to RBC lysis and staining. For intracellular staining, cells were fixed using BD Cytofix (BD Biosciences) and permeabilized using BD Phosflow Perm Buffer III according to manufacturer’s instructions. Staining of unfixed cells was performed at 4°C except for anti-CXCR5 Ab which was performed at room temperature. Stained cells were analyzed on a BD LSR II or BD FACSymphony A3 (BD Biosciences) using FACSDiva Software. Sorting experiments were performed on a BD ARIA SORP high-performance cell sorter (BD Biosciences). Flow cytometry data was analyzed using FlowJo Software version 10.8.1 (BD).

## KEY RESOURCES TABLE

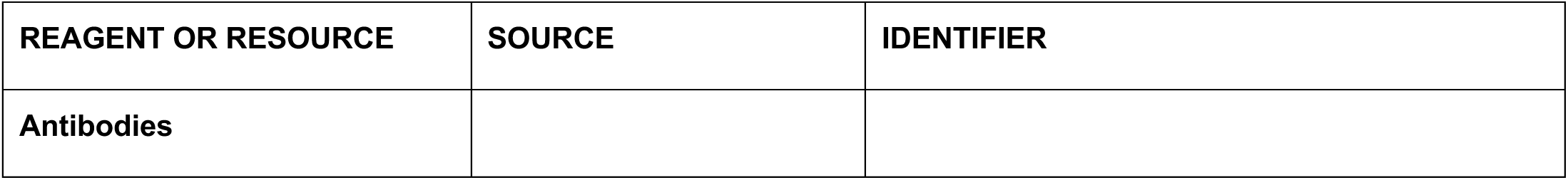
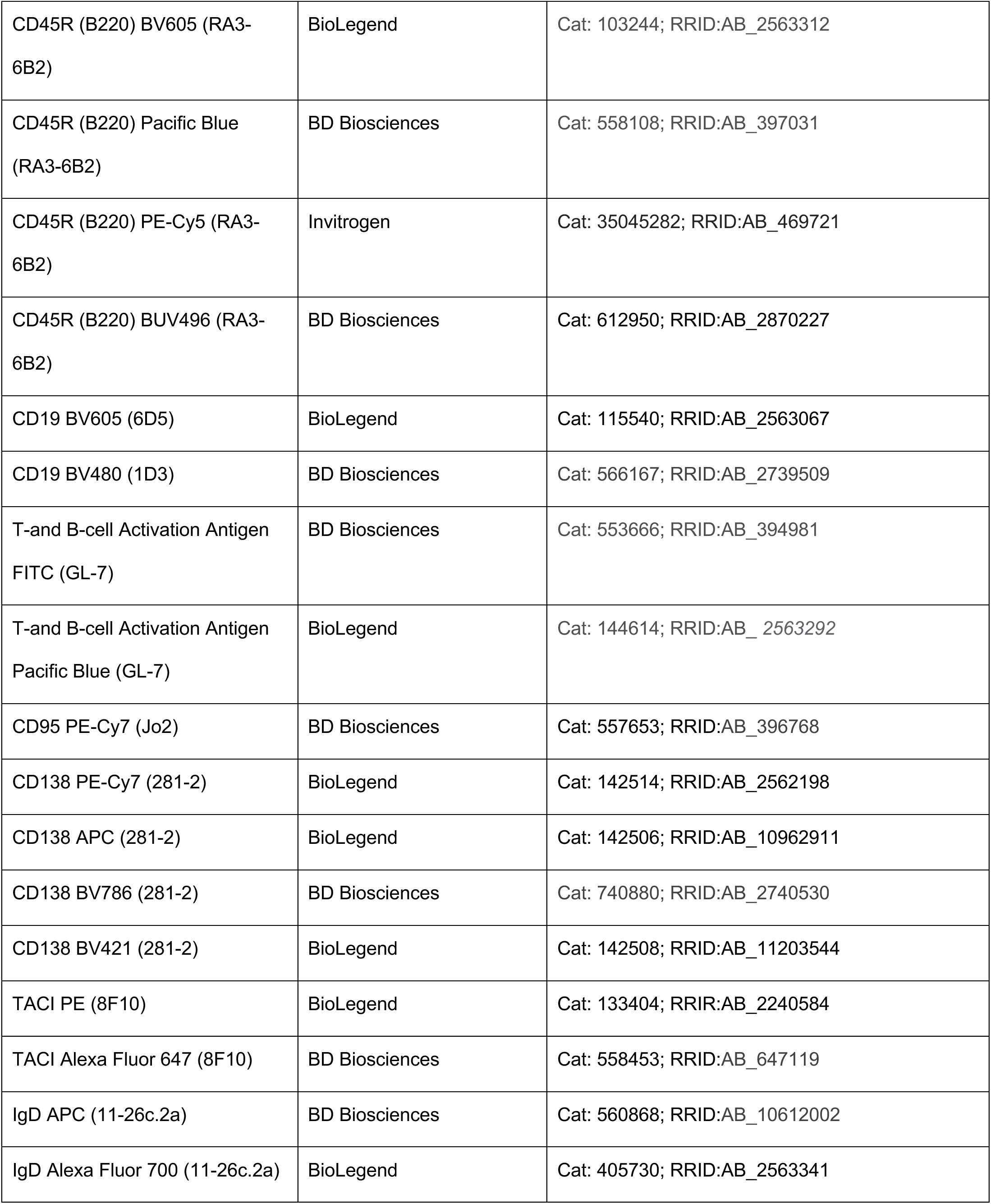
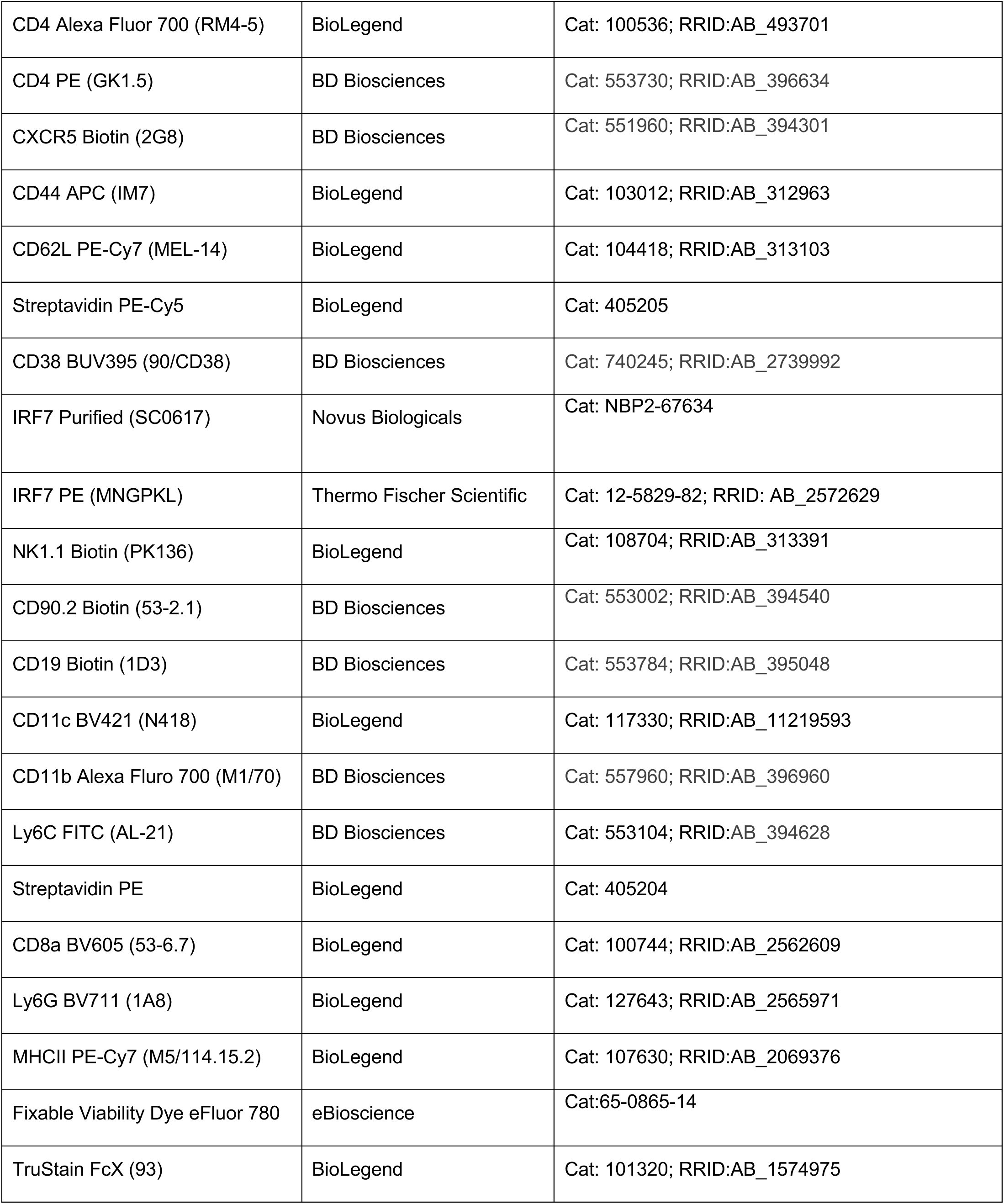
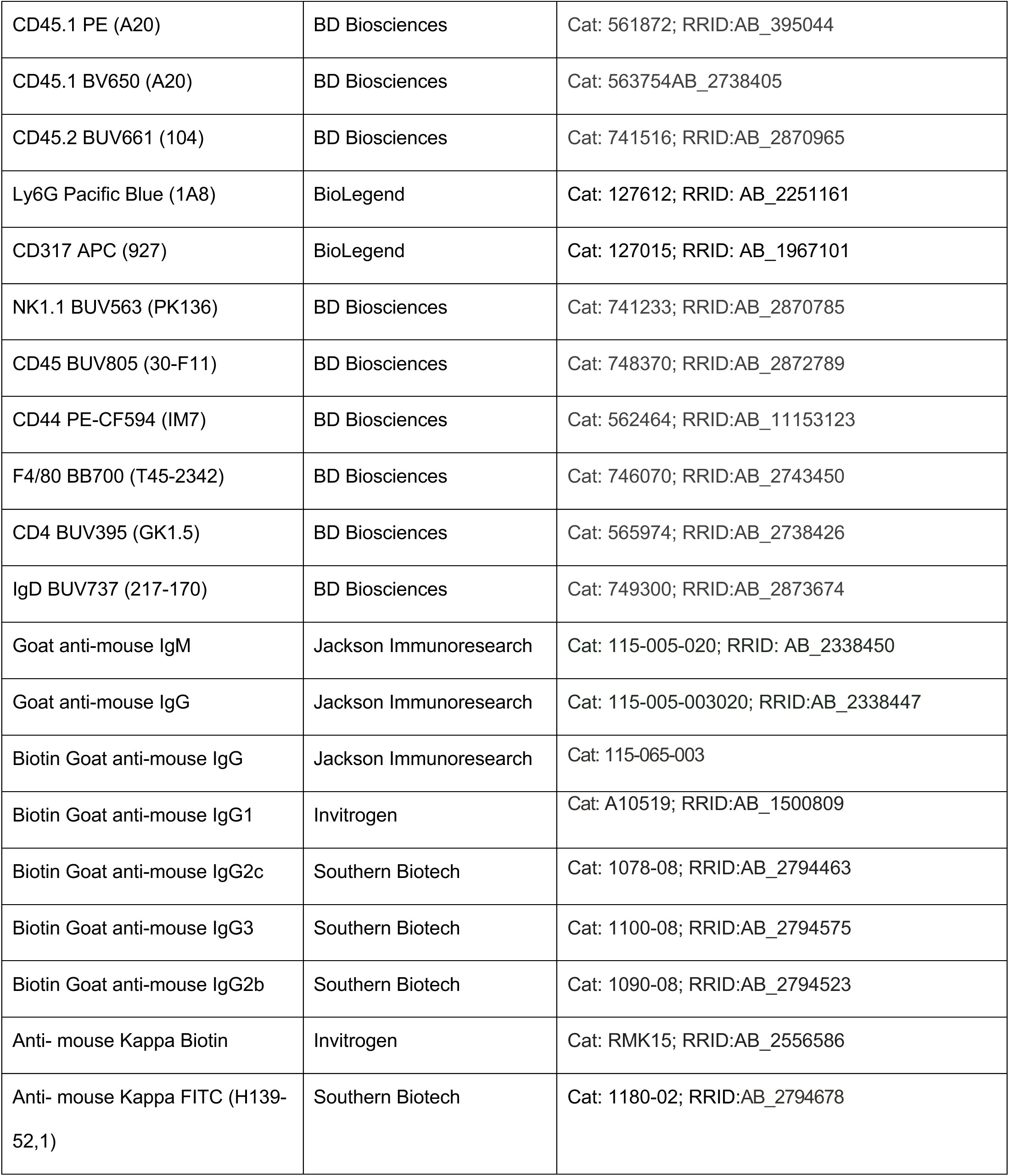
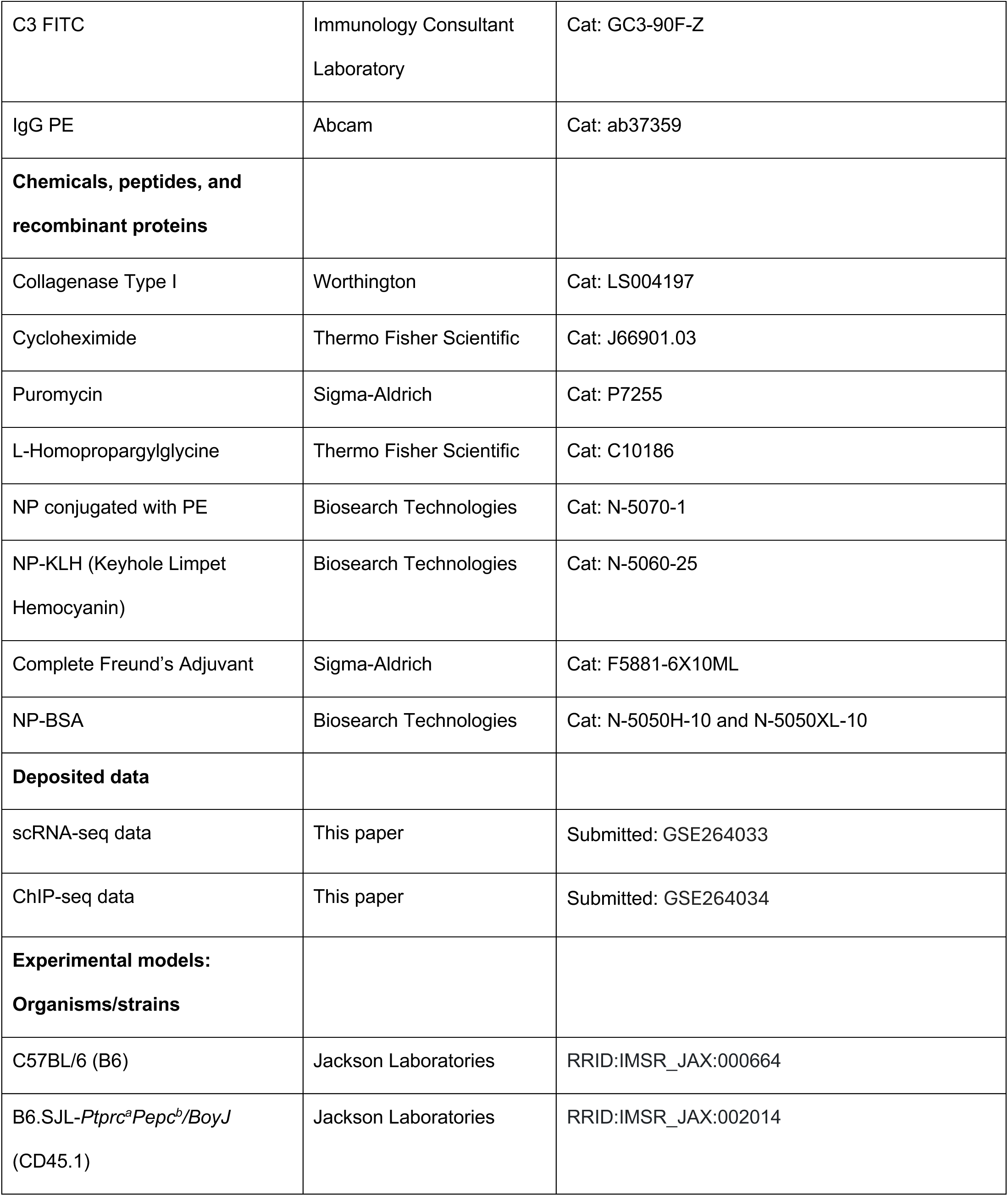
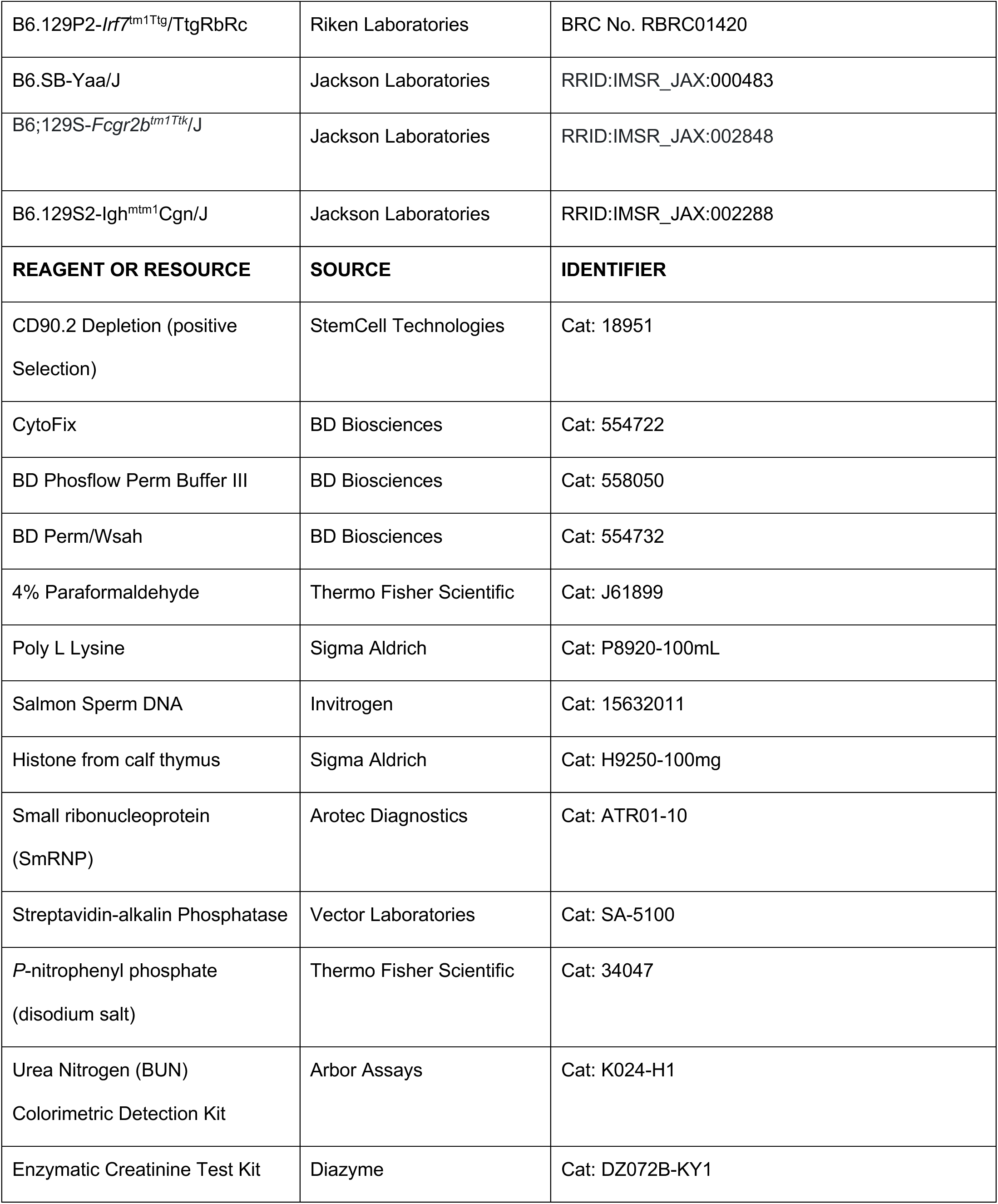
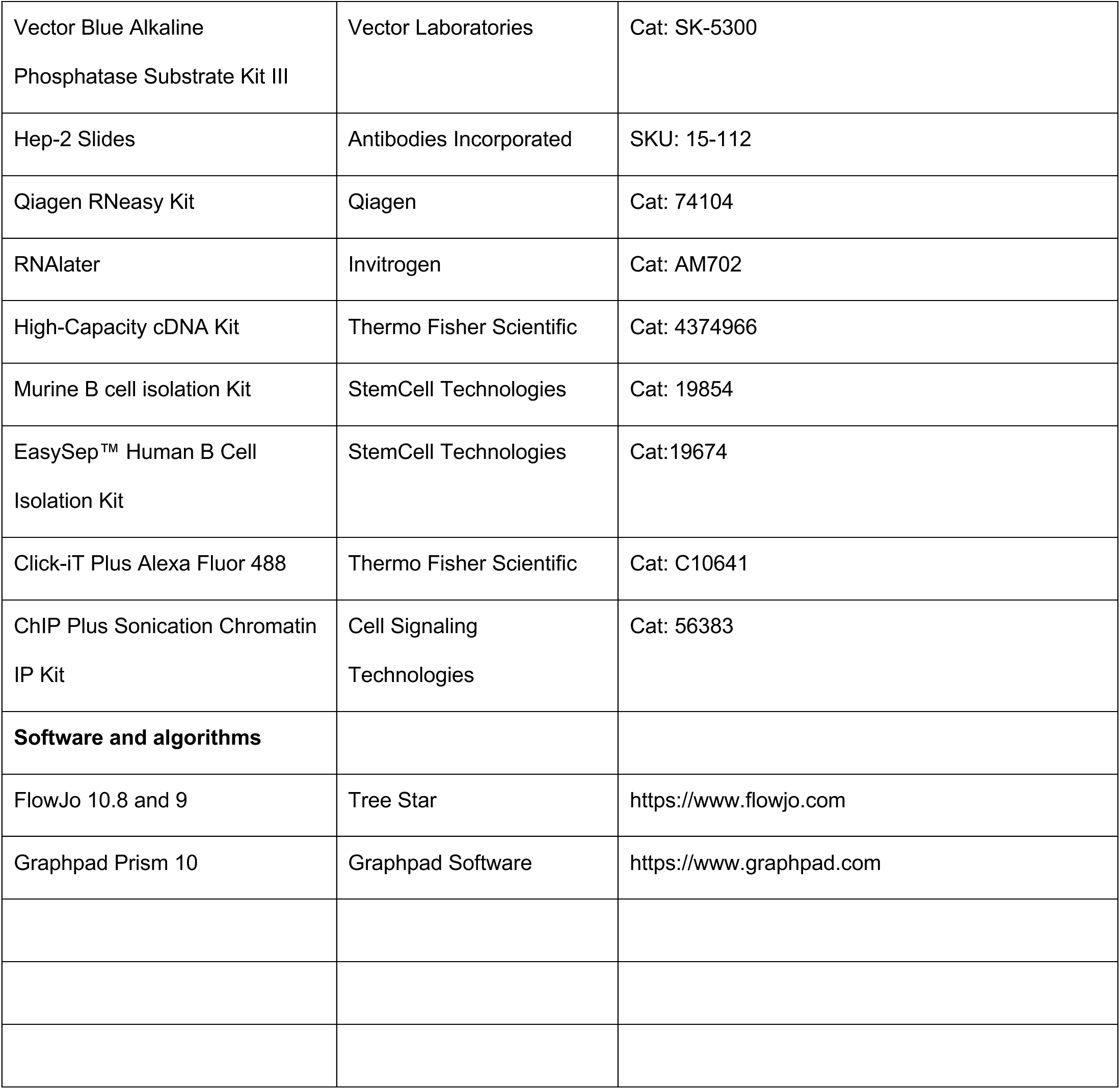

### Kidney histopathology and scoring

Kidneys from 6-mo-old mice were fixed in 10% neutral buffered formalin and embedded in paraffin. Kidney sections were cut at 3-μm thickness for periodic acid–Schiff. The slides were scanned on a TissueGnostics TissueFAXS SL imaging system with Kromnigon Spectra Split 7 filters and the TissueFAXS 71.1.138 software. Renal disease was evaluated with a semiquantitative 0-4+ scale as described (Chan et al., 1997). The scale was used for each compartment (glomerular, interstitial, and vascular) with the pathology graded according to specified criteria as absent (0-1), mild (1-2), moderate (2-3), or severe (3-4+). For comparative purposes, all the tissue sections were scored by one observer (R.C.), who was blinded to the origin of the kidney samples.

### ELISA

To detect serum antinuclear antibodies, ELISA plates (Thermo Fisher) were first coated with poly(L) Lysine followed by salmon sperm dsDNA (Invitrogen), nucleosome (histone from Sigma Aldrich on a layer of dsDNA coating), or small ribonucleoprotein (SmRNP) (Arotec Diagnostics). To detect NP-specific Abs, plates were coated with 10μg/mL NP4-BSA or NP29-BSA. Plates were washed and blocked with 5% FBS in PBS and serum was added starting at a 1:50 dilution in 5% FBS, followed by 1:2 serial dilution for the remainder of the plate. Antibodies were then detected with one of the following biotinylated antibodies: goat anti-mouse IgM (Jackson Immunoresearch Laboratories), goat anti-mouse IgG (Jackson Immunoresearch Laboratories, goat anti-mouse IgG2c (Southern Biotech). Plates were then washed and incubated with streptavidin-alkaline phosphatase (Vector Laboratories). *P*-nitrophenyl phosphate (disodium salt) (ThermoFisher Scientific) substrate for alkaline phosphatase was used for developing the plates and plates were read at λ405nm on Synergy H1 (BioTek Instruments). Serum BUN and Creatinine levels were measured via the Urea Nitrogen (BUN) Colorimetric Detection Kit (Arbor Assays) and the Enzymatic Creatinine Test Kit (Diazyme), respectively according to manufacturer’s instructions. All ELISA plates were read on Synergy H1 (BioTek Instruments) at the wavelengths indicated by the manufacturer’s instructions.

### ELISpot

For detection and quantification of autoantibody producing AFCs, multiscreen 96-well filtration ELISpot plates (Millipore Sigma) were coated with salmon sperm dsDNA (Invitrogen), nucleosome (histone from Sigma Aldrich on a layer of dsDNA coating), or Small ribonucleoprotein (SmRNP) (Arotec Diagnostics). To detect NP-specific Abs, ELISpot plates were coated with 10μg/mL NP4-BSA or NP29-BSA. Plates were blocked with 5% FBS in PBS. 2×10^6^ splenocytes suspended in RPMI 1640 (Corning) containing 10% FBS (Hyclone GE), Penicillin-streptomycin (Corning), and Glutamax (Corning) were added to the top row and serially diluted by twofold to the remainder of the rows. Plates were washed and autoantibody producing AFCs were detected by incubating with biotinylated anti-κ antibody (Invitrogen) followed by streptavidin-alkaline phosphatase (Vector Laboratories). Plates were then washed and developed via Vector Blue Alkaline Phosphatase Substrate Kit III (Vector Laboratories). ELISpot plates were analyzed using an Immunospot S6 Universal M2 (CTL).

### Immunofluorescence (IF) Microscopy and Hep-2 assay

For the detection of antinuclear antibody (ANA) seropositivity, serum from mice was diluted 1:50 and incubated on a Hep-2 slide (Antibodies Incorporated). ANAs were then detected using a FITC-rat anti-mouse k antibody (H139-52,1; Southern Biotech). ANA slides were visualized using a Leica DM4000 fluorescence software (Leica Microsystems).

Splenic and kidney IF microscopy was performed by embedding the tissue in OCT compound (Thermo Fisher Scientific) and snap freezing over liquid nitrogen. Five-µM sections were cut and mounted on a ColorFrost Plus Microscope Slide (Thermo Fisher Scientific) and fixed in cold acetone for 20 minutes. For splenic GC staining, tissue sections were stained with the following antibodies: GL-7 FITC and anti-IgD-APC (11-26c2a; BD Biosciences) and anti-CD4-PE (GK1.2; Biolegend). GC measurements were performed on a random selection of GL-7^+^ staining across the splenic tissue section where total GC area (µM^2^) was measured using the Leica Application Suite Advanced Fluorescence quantification tool. Kidney sections were stained with anti-CD3-FITC (Immunology Consultant Laboratory (ICL)) and anti-IgG-PE (Abcam) for the detection of IC depositions. Stained tissue sections were imaged on a Leica DM4000 Fluorescence microscope (Leica Microsystems).

### Seahorse Extracellular Flux Analysis

Splenic B cells were isolated from indicated mice via negative selection (StemCell Technologies). Cells were resuspended in XF DMEM pH 7.4 media supplemented with 1mM pyruvate, 2mM glutamine, and 10mM glucose and plated on poly-L-lysine coated microchamber wells. Oxygen consumption rates (OCR) and extracellular acidification rates were measured using a XF HS mini analyzer (Agilent). For the mito stress test, 1.5μM oligomycin, 2μM FCCP, and 0.5μM antimycin/rotenone were utilized. Data were analyzed using the Agilent Seahorse Wave Software.

### Translation Assay

Single cell suspension of splenocytes was prepared as described above. Splenocytes were incubated for 30 min in methionine-free RPMI 1640 medium with 10% of dialyzed FBS at 37°C. Cells were cultured for additional 2hr for HPG incorporation (final concentration 100 μM). Cycloheximide (100ug/ml) or puromycin (10ug/ml) treated samples were used as negative controls. HPG incorporation in GC B cells and plasma cells was stained with Click-iT Plus Alexa Fluor 488 Picolyl Azide Toolkit using Foxp3-fixation-permeabilization method and was detected by flow cytometry.

### Human B cell in vitro stimulation and IRF7 expression

PBMCs were isolated from two healthy donor whole blood samples using a Ficoll Gradient (BD) under an IRB-approved protocol at Penn State College of Medicine. B cells were then isolated from the PBMCs using StemCell EasySep Human B Cell Isolation Kit. Isolated B cells were stimulated overnight with 300ng/mL of R848 in 10% RPMI. RNA was isolated from B cells by the Qiagen RNeasy kit (Qiagen) according to manufacturer’s instructions. RNA quality and quantity were assessed via nanodrop prior to cDNA synthesis (Thermo Fisher Scientific). cDNA was generated using the High-Capacity cDNA kit (Thermo Fisher Scientific) according to manufacturer’s instructions. TaqMan primers for IRF7 (Hs01014809_g1) were used. Expression levels were normalized by a housekeeping gene (β-actin), and fold change calculated by the 2(ΔΔCt) method.

### ScRNAseq and Pathway Analysis

Spleens from two 3mo old FcγRIIB^−/−^ and two FcγRIIB^−/−^IRF7^−/−^ mice were processed for cell sorting as described above. CD3^−^B220^+^CD19^+^IgD^−^ and CD3^−^B220^+^CD19^+^IgD^+^ cells were sorted from each animal and mixed at a ratio of 9:1 (IgD^−^: IgD^+^) prior to library preparation. Library preparation and sequencing were performed by the Center for Applied Genomics Core at the Children’s Hospital of Philadelphia. Libraries were prepared using the 10x Genomics Chromium Single Cell 3’ Reagent kit v2 per manufacturer’s instructions. Sequencing was performed on an Illumina HiSeq sequencer. Data was then processed through the Cellranger pipeline (10x Genomics). Demultiplexing and alignments were performed against the mm10 transcriptome. Analysis was performed via Seurat package (v. 4.99) within R. Cells expressing less than 200 or more than 4000 unique genes were excluded from further analysis. Cells with >10% mitochondrial genes expression were also excluded. Log normalization and scaling of features was completed before principal component dimensionality reduction, clustering, and visualization via UMAP. Pseudotime analysis was performed using the Monocle3 package in R. A wilcoxon rank sum test with a Bonferroni multiple testing correction was used to identify differentially expressed genes between FcγRIIB^−/−^ and FcγRIIB^−/−^IRF7^−/−^ groups. Gene pathway-based analyses from the “Pre-GC transition” subclusters was performed using Ingenuity Pathway Analysis (IPA) Software (Qiagen).

### ChIPseq and analysis

Spleens from 10 five 4 month old FcγRIIB^−/−^ mice were pooled in duplicate (5 mice in each sample) and processed for cell sorting as described above. CD3^−^B220^+^CD19^+^IgD^−^ B cells were sorted and prepared for ChIP as previously described (Fike et al., 2023). Briefly, cells were fixed using 1.5% formaldehyde for 15 minutes at 37C followed by glycine to stop the reaction. Samples were processed further using Cell Signaling ChIP Plus Sonication Chromatin IP Kit (Cell Signaling) according to manufacturer’s instructions, Sonication was performed on a Biorupter PICO (Diagenode). Library prep and sequencing was performed by the Center for Applied Genomics Core at the Children’s Hospital of Philadelphia. Data was processed and analyzed in Bioconductor. Sequences were aligned to reference genome mm9. Peaks were called against input controls using Macs2. Peaks were visualized in IGV.

## QUANTIFICATION AND STATISTICAL ANALYSIS

All data were plotted as the mean of the data ± standard error of the mean (SEM), unless otherwise indicated. For all datasets, the normality of the data was first assessed based on the number of samples via D’Agostino-Pearson omnibus normality test or the Shapiro-Wilk normality test. Statistical significance between two groups was determined using the Student’s *t*-test when two groups were compared. For datasets containing more than two groups and multiple time points, a two-way ANOVA, with a Dunn-Sidak correction for multiple comparisons was performed. For the survival curve, a log-rank (Mantel-Cox) test was performed. The statistical test used is denoted in each figure legend. Statistical analyses were performed using Prism GraphPad software version 6 or 10 (GraphPad Software). P values are shown as NS, not significant *, p <0.05, **, p <0.01, ***, p <0.001, ****, p <0.0001 for significance.

## Acknowledgments

We would like to thank the Pennsylvania State University Hershey Medical Center flow cytometry core facility and Department of Comparative Medicine for help with flow cytometric experiments and animal housing and care, respectively. We thank the support staff within the central facility of the Department of Microbiology and Immunology at Pennsylvania State University College of Medicine. We would like to acknowledge the contributions of Dr. Renata Pellegrino Da Silva, Fernanda Thompson, Jacqueline Smiler, and the other members of the Center for Applied Genomics at CHOP for their help with the ScRNAseq and ChIPseq. We acknowledge the Sanderson Center for Optical Experimentation (SCOPE) and MLSC (The <u>SCOPE RRID</u> is SCR_022721) and Leah Whiteman for technical assistance. This project was partially funded under a grant with the Pennsylvania Department of Health using Tobacco CURE Funds, which specifically disclaims responsibility for any analyses, interpretations, or conclusions.

## Funding

National Institutes of Health grant RO1 AI162971 (ZSMR)

Finkelstein Memorial Award (AJF)

American Association of Immunology Careers in Immunology Fellowship (AJF)

National Cancer Institute 5T32CA060395-25 (AJF)

Lupus Foundation of America Goldie Simon Preceptorship Award (AJF)

National Heart Lung and Blood Institute (NHLBI) grant to MT: R35 HL150778

## Author contributions

Conceptualization: AJF, SBC, ZSMR

Methodology: AJF, ZSMR

Investigation: AJD, MVG, KNB, SBC, SME, JW, SAL, NMC

Formal Analysis: AJF, MVG, RC, RS, ZSMR

Funding acquisition: AJF, ZSMR

Resources: NJO, MT, DJL

Writing-original draft-AJF, ZSMR

Writing-review & editing-AJF, SBC, ZSMR

## Data and materials availability

“All data are available in the main text or the supplementary materials.” Single-cell-RNA-seq and ChIP-seq data have been deposited at the NCBI Gene Expression Omnibus and will be publicly available as of the date of publication (GSE264033 and GSE264034). Accession numbers are listed in the key resources table. Any additional information required to reanalyze the data reported in this paper will be available from the lead contact upon request.

