## Supplemental materials for "IRF7 controls spontaneous autoimmune germinal center and plasma cell checkpoints"

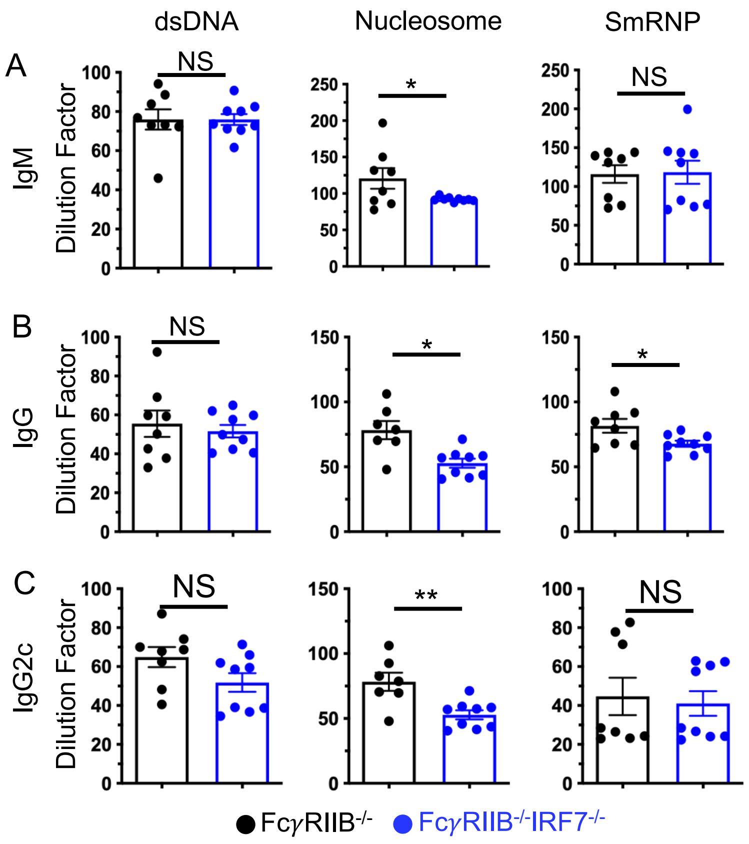


**Figure S1: Serum autoantibody titers at the early stages of systemic autoimmune responses.** dsDNA-, nucleosome- and Sm/RNP-specific serum IgM (**A**), IgG (**B**) and IgG2c (**C**) titers were measured in 2 month old Fc$\gamma$RIIB$^{-/-}$ and Fc$\gamma$RIIB$^{-/-}$IRF7$^{-/-}$ mice by ELISA. Each symbol represents an individual mouse and data are presented as means ± SEM. P values were calculated via an unpaired Student's t-test or Mann-Whitney test (NS, not significant, *, $p < 0.05$, **, $p < 0.01$.

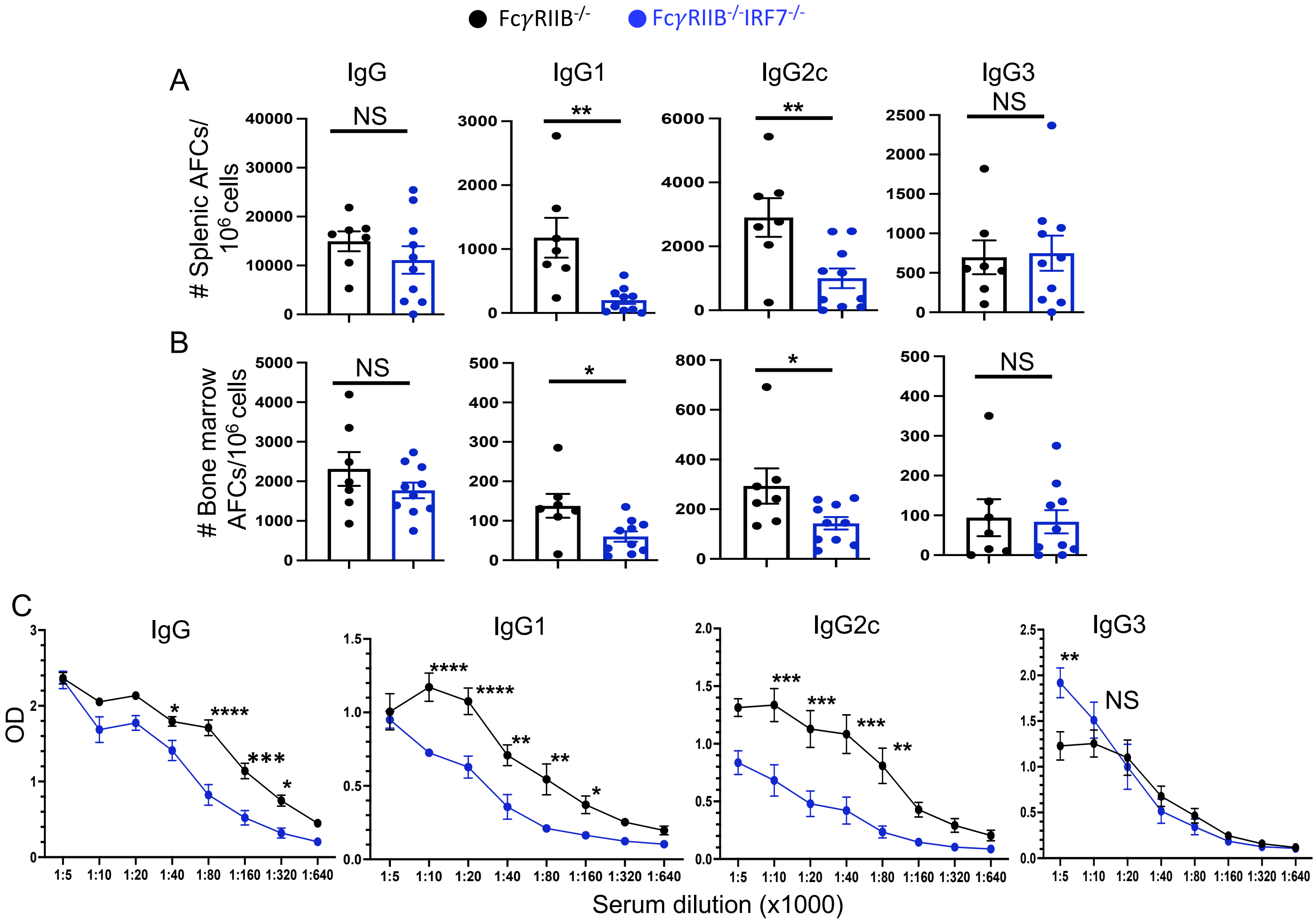


**Figure S2: Reduced numbers of steady-state IgG1- and IgG2c-producing AFCs and total serum IgG1 and IgG2c titers in the absence of IRF7. (A)** Numbers of splenic- and (**B**) Bone marrow AFCs were enumerated by ELISpot assay. (**C**) Total serum titers of IgG and IgG subclasses (IgG1, IgG2c and IgG3) in 4 month old Fc$\gamma$RIIB$^{-/-}$ and Fc$\gamma$RIIB$^{-/-}$IRF7$^{-/-}$ mice. Each symbol represents an individual mouse and data are presented as means ± SEM. P values were calculated via an unpaired Student's t-test or Mann-Whitney test (ns, not significant, *, p <0.05, **, p <0.01, ***, p <0.001). ****, p <0.0001.

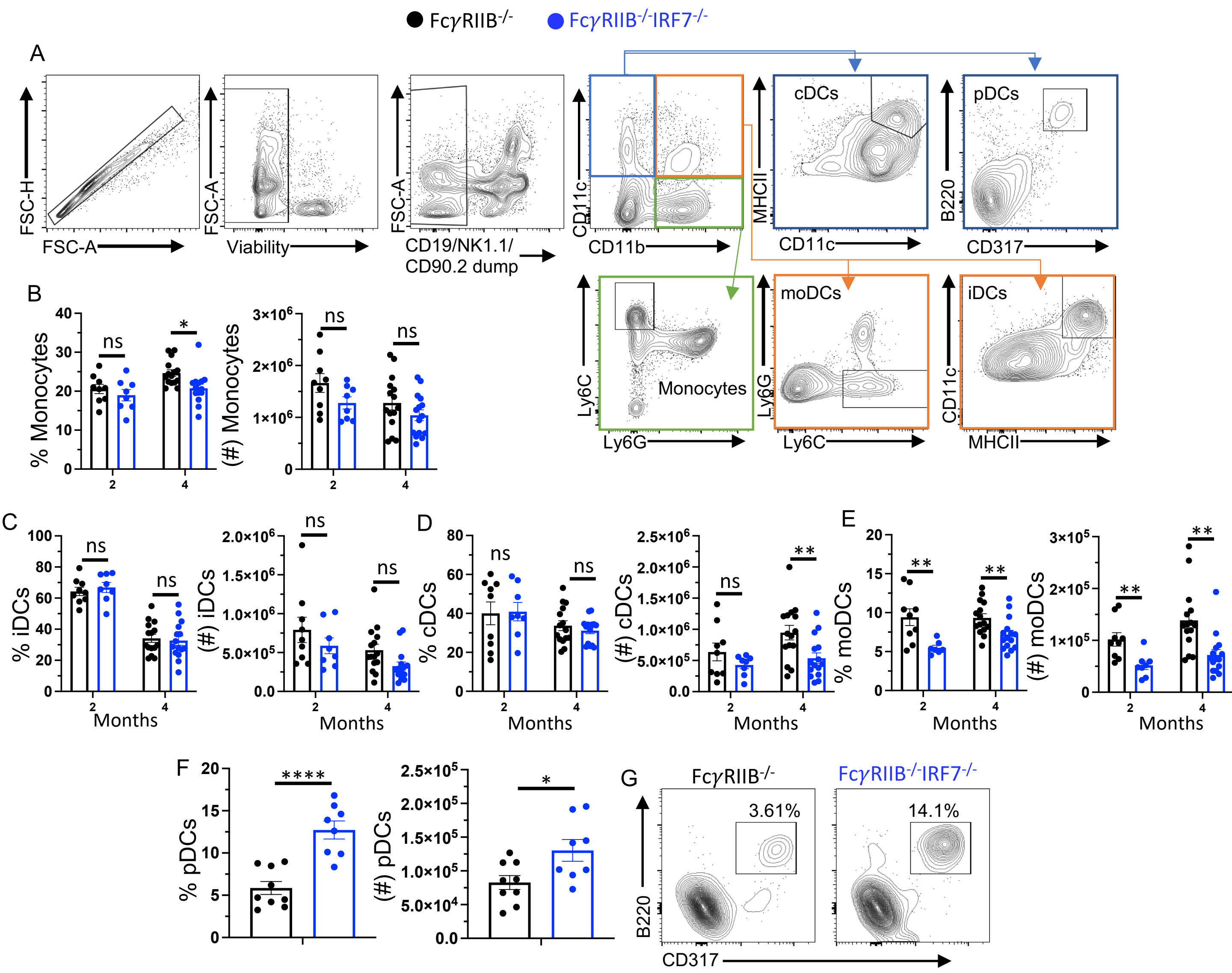


**Figure S3: Reduced number of cDCs and moDCs but increased number of pDCs in IRF7-deficient mice**. (A) The gating strategy of the flow cytometry data showing various myeloid cell populations such as conventional (cDCs), plasmacytoid (pDCs), monocyte derived (moDCs) and inflammatory (iDCs) DCs. Percentages and numbers of monocytes (B), iDCs (C) cDCs (D) moDCs (E) and pDCs (F, G) in 2 and 4 month old FcγRIIB$^{-/-}$ and FcγRIIB$^{-/-}$IRF7$^{-/-}$ mice are shown. (Nucleosome-, dsDNA- and SmRNP-specific serum IgM, IgG and IgG2c titers were measured in 4 and 6 month old FcγRIIB$^{-/-}$ and FcγRIIB$^{-/-}$IRF7$^{-/-}$ mice. Each symbol in (B-F) represents an individual mouse and data are presented as means ± SEM. P values were calculated via an unpaired Student's t-test or Mann-Whitney test (ns, not significant, *, p <0.05, **, p <0.01, ****, p <0.0001).

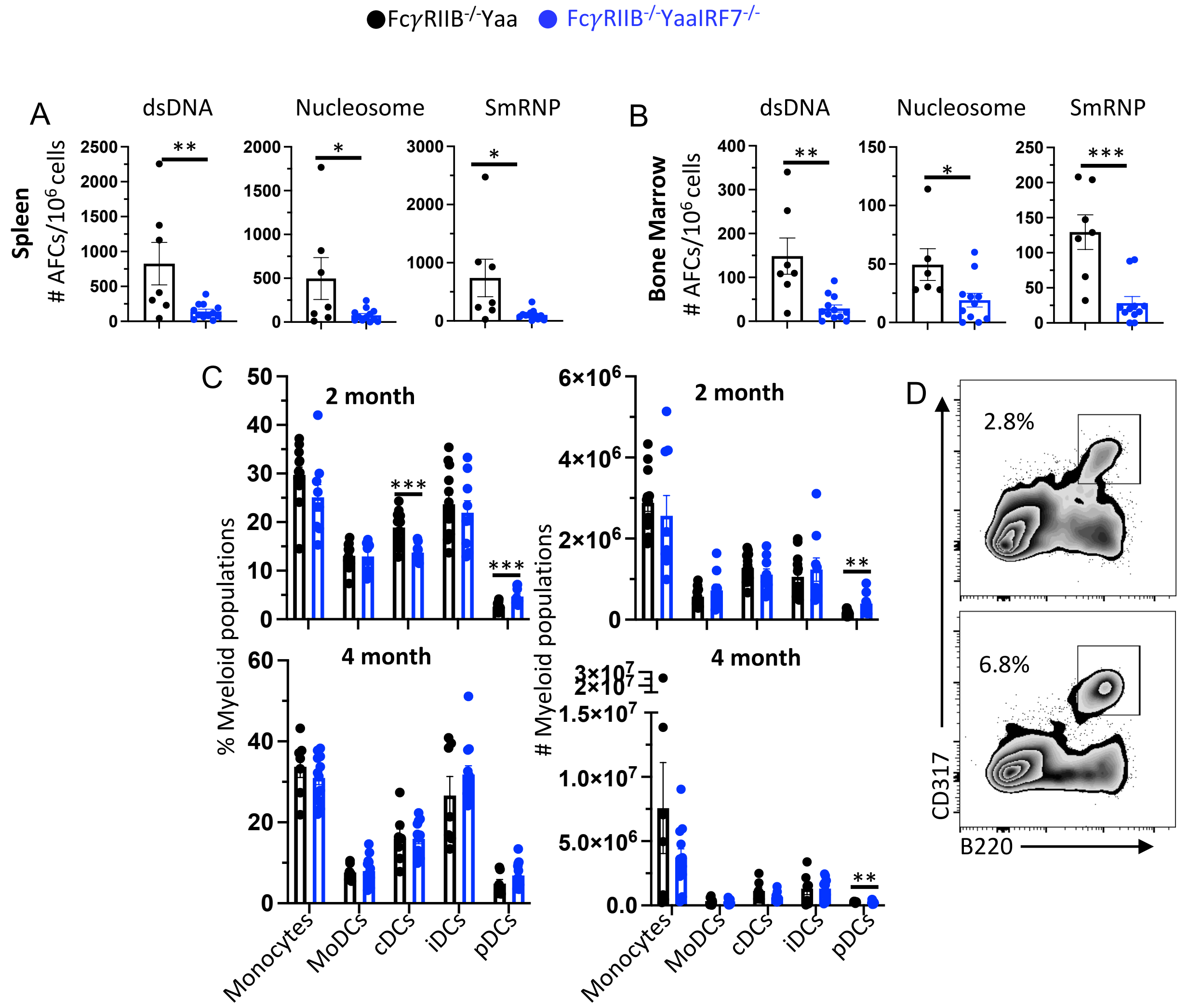


**Figure S4: Reduced autoreactive AFC and increased pDC numbers in Fc$\gamma$RIIB$^{-/-}$YaaIRF7$^{-/-}$ mice.** The number of dsDNA-, nucleosome-, and SmRNP-specific splenic (**A**) and bone marrow (**B**) AFCs was quantified by ELISpot in 3 month old Fc$\gamma$RIIB$^{-/-}$ Yaa and Fc$\gamma$RIIB$^{-/-}$ YaaIRF7$^{-/-}$ mice. (**C**) Percentage and number of myeloid cells in these mice. (**D**) Gating strategy of B220$^{+}$CD317$^{+}$ (PDCA-1) pDCs. Each symbol represents an individual mouse and data are presented as means ± SEM. P values were calculated via an unpaired Student's t-test or Mann-Whitney test (ns, not significant, *, p <0.05, **, p <0.01, ***, p <0.001).

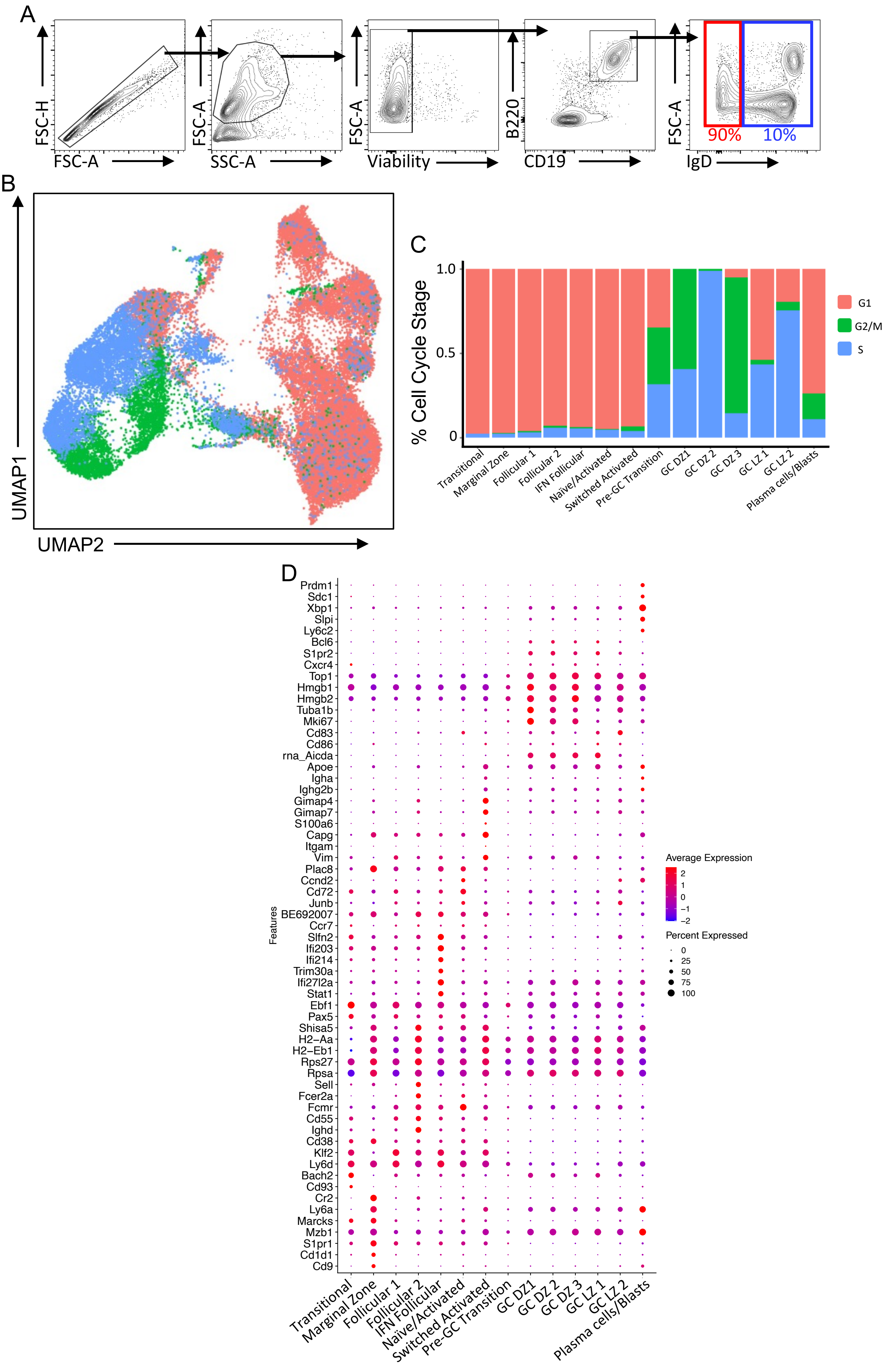
A
FSC-H
FSC-A
FSC-A
SSC-A
FSC-A
Viability
B220
CD19
FSC-A
IgD
90%
10%
B
UMAP1
UMAP2
C
% Cell Cycle Stage
1.0
0.5
0
G1
G2/M
S
Transitional
Marginal Zone
Follicular 1
Follicular 2
IFN Follicular
Naïve/Activated
Switched Activated
Pre-GC Transition
GC DZ1
GC DZ 2
GC DZ 3
GC LZ 1
GC LZ 2
Plasma cells/Blasts
D
Features
Prdm1
Sdc1
Xbp1
Slpi
Ly6c2
Bcl6
S1pr2
Cxcr4
Top1
Hmgb1
Hmgb2
Tuba1b
Mki67
Cd83
Cd86
rna_Aicda
Apoe
Igha
Ighg2b
Gimap4
Gimap7
S100a6
Capg
Itgam
Vim
Plac8
Ccnd2
Cd72
Junb
BE692007
Ccr7
Slfn2
Ifi203
Ifi214
Trim30a
Ifi27l2a
Stat1
Ebf1
Pax5
Shisa5
H2−Aa
H2−Eb1
Rps27
Rpsa
Sell
Fcer2a
Fcmr
Cd55
Ighd
Cd38
Klf2
Ly6d
Bach2
Cd93
Cr2
Ly6a
Marcks
Mzb1
S1pr1
Cd1d1
Cd9
Average Expression
2
1
0
−1
−2
Percent Expressed
0
25
50
75
100
Transitional
Marginal Zone
Follicular 1
Follicular 2
IFN Follicular
Naïve/Activated
Switched Activated
Pre-GC Transition
GC DZ1
GC DZ 2
GC DZ 3
GC LZ 1
GC LZ 2
Plasma cells/Blasts

**Figure S5. Single cell transcriptome of autoimmune-prone B cells identifies several clusters**. (**A**) Gating strategy for sorting 90% $IgD^-$ activated and 10% $IgD^+$ naïve B cells for scRNAseq analysis. (**B**) UMAP depicting cell cycle status of clusters determined based on average gene expression of gene sets annotated for each cell cycle state. (**C**) Percentage of each cell cycle phase per cluster. (**D**) Dot plot depicting some of the major markers used in the annotation of the scRNAseq of B cell clusters where dot color indicates the average expression of the gene within that cluster and the size of the dot indicates the percentage of the cluster expressing that gene.
